# Maximum Antigen Diversification in a Lyme Bacterial Population and Evolutionary Strategies to Overcome Pathogen Diversity

**DOI:** 10.1101/2020.12.16.423180

**Authors:** Lia Di, Saymon Akther, Edgaras Bezrucenkovas, Larisa Ivanova, Brian Sulkow, Bing Wu, Maria Gomes-Solecki, Wei-Gang Qiu

## Abstract

Natural populations of microbes and their hosts are engaged in an arms race in which microbes diversify to escape host immunity while hosts evolve novel immunity. This co-evolutionary process, known as the “Red Queen” hypothesis, poses a fundamental challenge to the development of broadly effective vaccines and diagnostics against a diversifying pathogen. Based on surveys of natural allele frequencies and experimental immunization of mice, we show minimal antigenic cross-reactivity among natural variants of the outer surface protein C (OspC), a dominant antigen of a Lyme Disease-causing bacterium (*Borrelia burgdorferi*). To overcome the challenge of OspC antigenic diversity to clinical development of preventive measures, we implemented a number of evolution-based strategies to broaden OspC immunological cross-reactivity. In particular, the centroid algorithm – a genetic algorithm to minimize sequence differences with natural variants – generated synthetic OspC analogs with the greatest promise as diagnostic and vaccine candidates against diverse Lyme pathogen strains coexisting in the Northeast United States. Mechanistically, we propose a model of runaway maximum antigen diversification (MAD) mediated by amino-acid variations distributed across hypervariable regions on the OspC molecule. Under the MAD model, evolutionary centroids display high cross-reactivity by occupying the central void in the antigenic space excavated by diversifying natural variants. In contrast to the vaccine design based on concatenated epitopes, the centroid algorithm generates analogs of native antigens and is automated. The MAD model and evolution-inspired antigen designs have broad implications for combating diversifying pathogens driven by pathogen-host coevolution.

**Importance:** Microbial pathogens rely on molecular diversity of cell surface antigens to escape host immunity. Vaccines based on one antigen variant often fail to protect the host against pathogens carrying other variants. Here we show evolution-based designs of synthetic antigens that are broadly reactive to all natural variants. The evolutionary analogs of a major surface antigen of a Lyme disease bacterium (*Borrelia burgdorferi*) showed promise as vaccine candidates against diverse pathogen strains coexisting in the endemic areas of Lyme disease in Northeast United States. Our evolution-based computational design is automated, generates molecular analogs of natural antigens, and opens a novel path to combating fast-evolving microbial pathogens.

## Introduction

### Rapid antigen evolution driven by “Red Queen” coevolution

Colloquially known as the “kill the winner” selection or the “Red Queen” hypothesis, negative frequency-dependent selection (NFDS) is an evolutionary mechanism that favors rare phenotypes over common ones, promoting biological novelty [1–3]. Driven by NFDS, antigenic variation is a host defense strategy ubiquitously shared among viral, bacterial, and eukaryotic pathogens [4–6]. Consequently, the power of NFDS in driving pathogen diversity becomes a fundamental challenge for developing broadly effective clinical measures against fast-evolving microbial pathogens especially viruses [7–9]. Although bacterial pathogens do not evolve as rapidly as viral pathogens, development of broad-spectrum anti-bacterial interventions is nonetheless hampered by a large number of cell-surface antigens encoded on bacterial genomes as well as by the vast allelic diversity segregating at antigen loci in natural pathogen populations [5].

Here we hypothesize that microbial pathogens and their surface antigens are under evolutionary constraints despite a trend of relentless diversification. Specifically, we propose and test the presence of an evolutionary constraint to antigen evolution as a result of NFDS – maximum antigenic diversification (MAD), the expectation that coexisting pathogen variants are obligatorily distinct from each other in antigenicity. The MAD hypothesis is a corollary of the strain theory of pathogen-host co-evolution, which posits that host immunity drives and molds pathogen populations into recognizable genetic units [3,6]. Under the strain model, coexisting microbial strains occupy high-fitness peaks on an antigenic landscape shaped by host immunity where any off-peak antigen variants (e.g., a recombinant variant) are at a selective disadvantage and would be eliminated by host immunity [3]. As such, the requirement for microbial genomes to perch on narrowly defined fitness peaks represents a major evolutionary constraint imposed by host immunity. This evolutionary constraint could be exploited to tip the balance of the “Red Queen” coevolution for the benefit of the host. For example, the precarious coexistence of pathogen strains could be destabilized and the pathogen population be driven to extinction if the host immunological landscape is re-molded by, e.g., an introduction of novel antigen variants as vaccines.

### Antigenic variations in Lyme disease pathogens

For over three decades, Lyme disease has been the most prevalent vector-borne disease in the United States and Europe [10]. It is caused by spirochetes of the *Borrelia burgdorferi sensu lato* species complex, also known as a new bacterial genus *Borreliella* [11,12]. A single species, *B. burgdorferi*, transmitted by *Ixodes scapularis* ticks in the Northeast and Midwest and *I. pacificus* in the West, causes the majority of Lyme disease cases in the US. Genes encoding cell surface lipoproteins are over-represented in the ~1.5 Mbp genome of *B. burgdorferi*, totaling 4.9% of the chromosomal genes and 14.5% of the plasmid-borne genes, in contrast to ~2.0% lip-oprotein-encoding genes in other bacterial pathogens such as *Helicobacter pylori* and *Treponema pallidum* [13]. Genome comparisons further revealed that lipoprotein-encoding genes are the most variable loci within the genome, consistent with their roles in evading vector and host immunity [14]. For example, *B. burgdorferi* shifts its surface lipoprotein composition when migrating between the tick vector and the mammalian host, up-regulating the expression of outer surface protein A (OspA) during tick colonization, the expression of OspC during host invasion, and the expression of VlsE (Variable membrane protein-Like Sequence E) during persistency within the host [15,16].

Among the large repertoire of genes encoding cell surface lipoproteins, *ospC* plays an outsized role in *B. burgdorferi* immune escape. First, *ospC* is required for initial invasion into the hosts, suggesting its role in defense against host innate rather than adaptive immunity [17,18]. Molecular functions of OspC include anti-phagocytosis and plasminogen-binding activities, both of which serve to abort host innate immune responses [19,20].

Second, OspC is an immuno-dominant and serotype-determinant antigen of *Borreliella* strains [21,22]. Experimental immunization of mice with recombinant OspC variants elicited strain-specific protective immunity against strains expressing homologous but not heterologous OspC variants [23,24]. Further, experimental immunization of mice using whole sera from infected mice showed that polyclonal antibodies binding OspC were the main components of strain-specific immunity [25]. Field-based studies further supported NFDS acting on the *ospC* locus being the main evolutionary mechanism maintaining genomic diversity in natural *B. burgdorferi* populations [26–29].

Third, sequence variations at *ospC* are in nearly complete linkage disequilibrium with genomic lineages in North America, suggesting within-population lineage diversification driven by *ospC* variability [27,30]. Simulations based on principles of population genetics showed that the nearly one-to-one correspondence between the major *ospC* sequence alleles and the co-existing *B. burgdorferi* lineages was consistent with a history of within-population genome diversification driven by NFDS targeting the *ospC* locus [27]. Additional evidence supporting the *ospC-*driven diversification of local *B. burgdorferi* lineages includes the high recombination rate at *ospC* and the uniform distributions of *ospC* alleles and genomic groups [29,31]. While it remains a possibility that sequence variation at *ospC* is driven by host diversity in this generalist parasite [32], the “multiple-niche” hypothesis appears to be inconsistent with results of field studies of *B. burgdorferi* populations in North America and *B. afzelii* populations in Europe as well as with results of direct experimental tests [33,34].

### Quest for broadly cross-reactive OspC molecules

Immuno-dominance of OspC makes it a valuable target for anti-Lyme diagnostics and vaccines, yet its clinical potentials are hindered by its sequence hyper-variability. Thus far, strategies to overcome OspC diversity have been focused on identifying conserved epitopes or a combination of epitopes shared among natural variants [35–37]. For example, we identified a minimum set of OspC variants as broadly effective diagnostics by measuring antigenicity of recombinant OspC variants using sera from immunized mice as well as sera from naturally infected hosts including the white-footed mice (*Peromyscus leucopus*), a reservoir species of *B. burgdorferi* in the Northeast US [37]. Conserved structural domains on the dimeric OspC molecule proved to be ineffective targets of vaccination [38]. A concatenation of eight OspC epitopes became the base of a broadly immunogenic vaccine for canine use [35,39]. In another systematic effort, protein arrays made of immobilized recombinant OspC variants were used to map key OspC epitopes to the hypervariable C-terminal region [36].

The MAD hypothesis suggests an alternative and novel strategy to overcome OspC diversity based on the Red Queen co-evolution. First, based on frequencies of antigen variants in nature as well as experimental immunization of mice, we tested maximum antigenic diversification among the 16 OspC variants coexisting in natural populations of the Lyme disease pathogens in the Northeast United States [27,40]. Second, we used evolutionary algorithms to design analogs of natural OspC molecules with minimal sequence differences to natural variants. We cloned and purified these synthetic OspC molecules and tested their antigenic breadth using sera from artificially and naturally infected hosts. Third, we explored molecular mechanisms underlying the broad antigenicity of evolutionary antigens with computer simulations. One of our evolution-based designs – the consensus algorithm – is similar to the COBRA approach used to design broadly reactive vaccines against the influenza virus [8]. Critically, the evolution-based strategies to overcome microbial diversity are automated and able to generate synthetic analogs that preserve the structure and function of natural antigen variants while by-passing the evolutionary constraints imposed on natural variants by host immunity.

## Results

### Lack of immune cross-protection among *B. burgdorferi* strains in nature

Previously, we used high-throughput deep sequencing of the *ospC* locus to quantify *B. burgdorferi* strain diversity within single *I. scapularis* ticks [41]. Consistent with earlier results based on DNA cloning and DNA-DNA hybridization, the results re-affirmed a largely uniform distribution of a diverse set of *B. burgdorferi* strains identifiable by ~16 major-group *ospC* alleles in the highly endemic regions of Lyme disease in the Northeast US [26,29]. Using the same dataset (Supporting Information S3a Data), in this study we tested if frequencies of pairs of *B. burgdorferi* strains co-infecting a single tick were higher, lower, or equivalent relative to the expectation of random allelic association. With a sample of *n*=119 infected *I. scapularis* ticks, we found that the majority of strain pairs were over-represented relative to random expectations and no pair was significantly under-represented in infected ticks (Fig 1). These results suggest a lack of cross-protective immunity in the reservoir hosts against infection by multiple *B. burgdorferi* strains in nature. Indeed, infection by one *B. burgdorferi* strain facilitates super-infection by additional strains [31,34]. We conclude that *B. burgdorferi* strains marked by OspC variants are antigenically distinct in the wild as has been shown experimentally [23,24].

**Fig 1.**
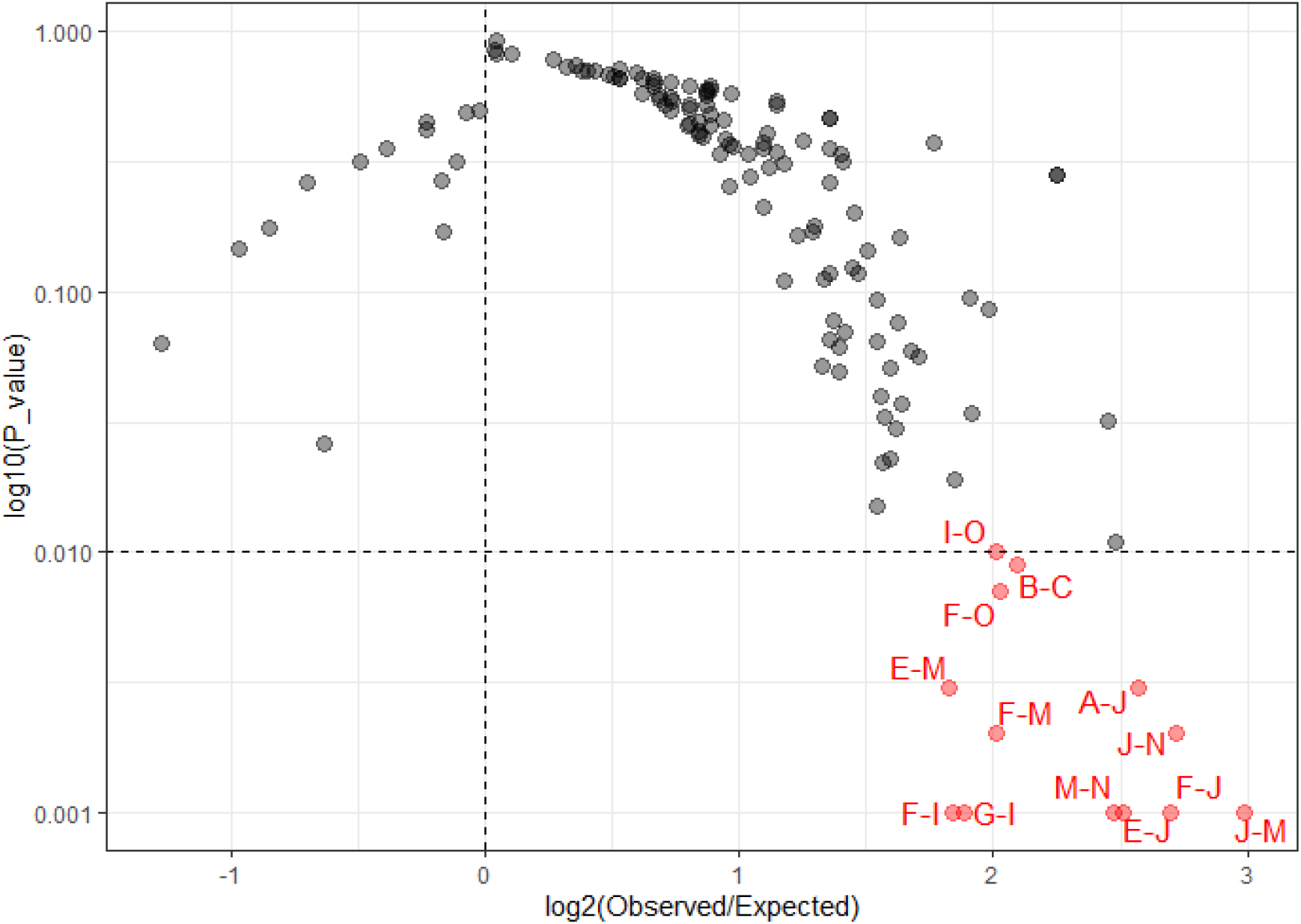
Over-representation of mixed *B. burgdorferi* strains infecting individual ticks. Each point represents a pair of OspC alleles detected in *N*=119 infected ticks by deep sequencing [31]. The co-occurrence of an OspC pair was quantified for the degree of over- or under-representation based on observed and expected frequencies (*x*-axis) and for statistical significance using random permutations of alleles among ticks (*y*-axis). Most OspC allelic pairs were more abundant than expected by chance (i.e., fold change > 0, with significant pairs at *p*<0.01 in red), suggesting weak immune cross-protection of reservoir hosts against co-infection by multiple *B. burgdorferi* strains in nature.

### Antigenic specificity of OspC variants tested with immunized C3H mouse sera using ELISA

We immunized the C3H mice with 16 recombinant OspC natural variants and tested their cross-reactivity based on binding with the OspC variant-specific sera using ELISA [36]. We identified six OspC variants (A, B, E, F, I, and K) most broadly reactive with the variant-specific sera, consistent with results using naturally infected sera from human patients, dogs, and *P. leucopus* mice [36]. Here we re-analyzed the ELISA dataset (Supporting Information S3b Data) by correcting for serum-to-serum variation (see Material & Methods).

The serum-normalized ELISA readings showed that, with two exceptions, rOspCs reacted significantly (i.e., with *z~*2.0, two standard deviations above the mean) with homologous sera, indicating high antigenic specificity of rOspCs (Fig 2, bar plot). The two exceptions included the variant F, which reacted significantly with both the F− and the B-specific sera, and the variant J, which reacted more strongly with the M-specific serum than with the J-specific serum. The high antigenic specificity of rOspCs is alternatively visualized with a heat map, which shows a strong diagonal line indicating the highest reactivity of rOspCs with homologous sera (Fig 2, heat map). Note an absence of L-specific sera in both the bar plot and the heat map. Note also that although heterologous bindings were weaker than homologous bindings, rOspCs typically nonetheless reacted with heterologous sera. In other words, a binding value of *z=*0 represents the normalized average binding level, not an absence of antigen-serum reaction.

**Fig 2.**
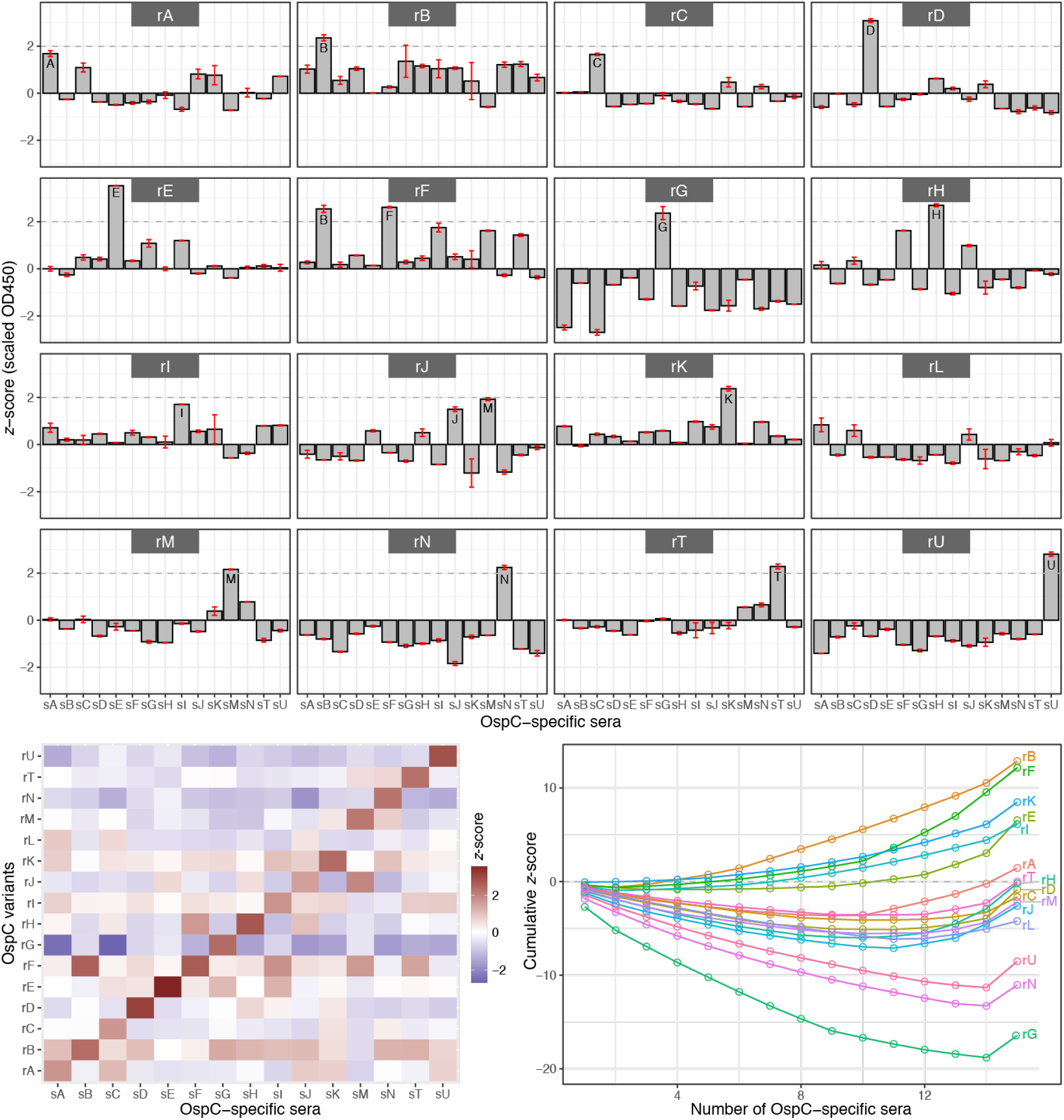
ELISA testing of C3H mice immunized with OspC variants. Fifteen sera (“sA” through “sU”) of mice, each immunized with a specific recombinant OspC variant, were assayed for reactivity with 16 OspC variants (“rA” through “rU”) using ELISA [37]. (*Top*) Each panel shows binding intensities (*y–*axis) of an OspC variant with a panel of OspC-specific sera (*x–*axis). Error bars show one standard deviation above and below the mean of two replicated assays. OD450 readings of each sera were normalized to a mean a zero and to the unit of standard error (i.e., *z–*score). A value above the *z*=2 line (dashes) indicates a highly significant reaction. (*Bottom left*) A heat map representation of mean *z*-scores of ELISA. (*Bottom right*) Antigen reaction characteristics (ARC) curves, similar to the receiver-operation characteristics (ROC) curves, is a measure of antigen specificity. Each curve traces cumulative *z–*scores (*y–*axis) of an OspC variant’s binding intensities with the sera samples, ordered by the lowest to the highest reactivity. The ARC curve rises with an above-average binding value (*z>*0) and drops with a below-average binding value (*z*<0). Thus, a high-rising curve (e.g., for rB) indicates consistently above-average reactivity with sera samples, suggesting a broadly cross-reactive antigen. Conversely, a low-lying curve (e.g., for rG) indicates consistently below-average reactivity, suggesting a relatively specific antigen. Curves close to the zero line (the majority of variants) indicate antigens with an average level of cross-reactivity.

In more quantitative details, an antigen reaction characteristic (ARC) curve captures the full spectrum of a rOspC’s antigenicity by showing cumulative binding levels with all serum samples (Fig 2, ARC curves). In addition, the ARC curves provide a quantitative measure of antigen specificity and cross-reactivity, showing the most broadly reactive antigens at the top and the most specific antigens at the bottom. For example, the ARC curves show B, F, K, E, and I being the most broadly reactive variants, highlighting the conclusion of the original study [36].

### Antigenic specificity of OspC variants tested with immunized C3H and *P. leucopus* mouse sera using immunoblots

We further tested the antigenic specificity of rOspCs using a full set of 16 variant-specific sera (the L-specific sera included) from immunized C3H and *P. leucopus* mice with immunoblot assays (Fig 3, top). The raw immunoblot images showed strong specific reactions of rOspC with homologous sera (diagonal) and weak reactions with heterologous sera (off-diagonal). As in the ELISA data analysis, we corrected for serum-to-serum variation by normalizing binding intensities for each serum (Supporting Information S3c Data). Consistent with ELISA results, the re-scaled intensities showed the strongest bindings between rOspCs with homologous sera (Fig 3, heat map). However, the ARC curves showed a lack of consistency in the topmost cross-reactive rOspC variants between the immunoblot using the C3H mice (rH and rI at the top) and the immunoblot using the *P. leucopus* mice (rT and rJ at the top) (Fig 3, ARC curves). Further, the rOspC rankings of the immunological breadth as quantified by the ARC curves in both immunoblots were different from the ranking from the ELISA experiment using the C3H mice (rB and rF at the top, Fig 2, ARC curves).

**Fig 3.**
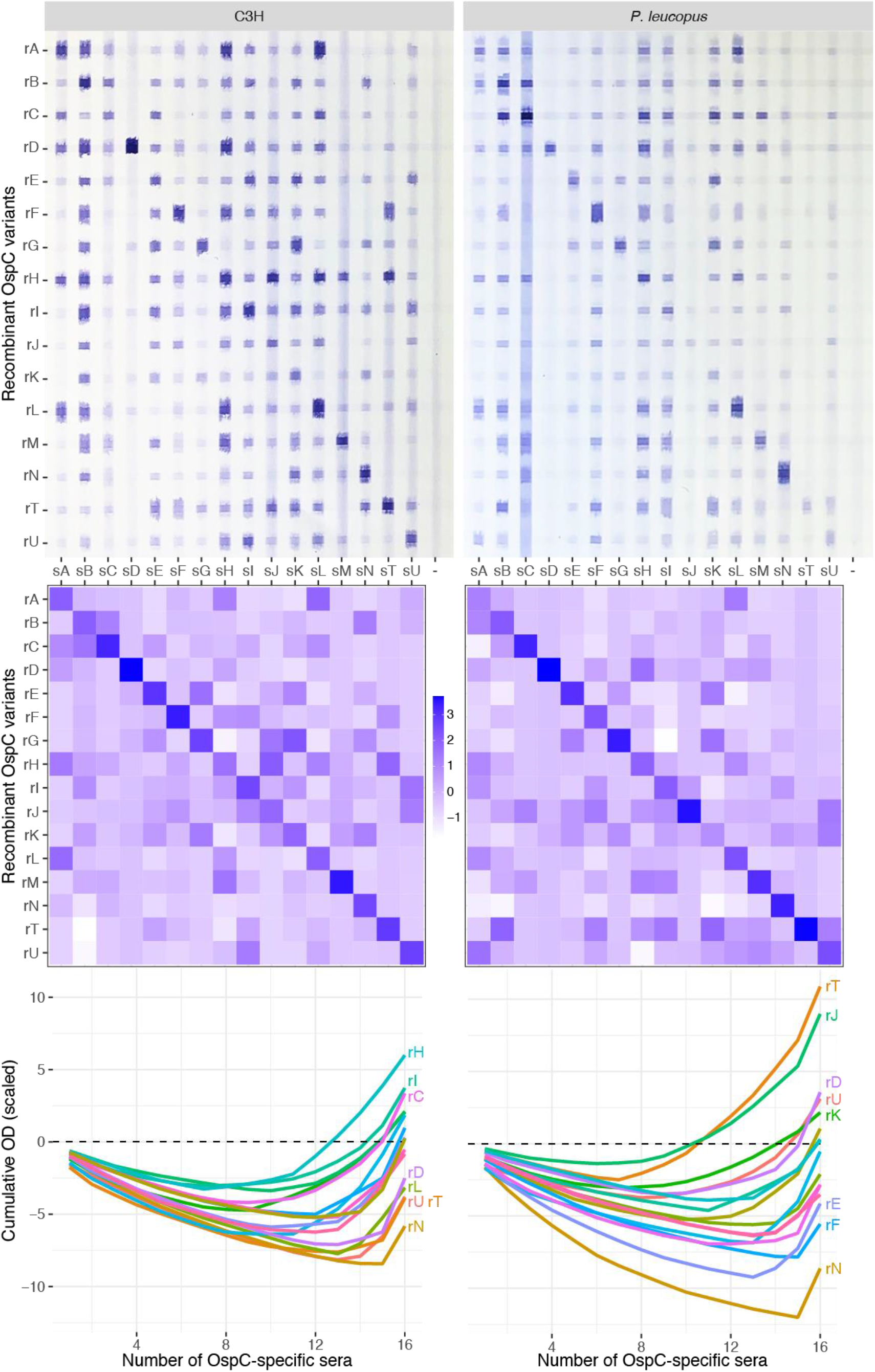
Immunoblot testing of C3H and P. leucopus mice immunized with OspC variants. (*Top*) Immunoblot images of OspC-specific sera (*x*–axis) from the C3H mice (*left*) and the *P. leucopus* mice (*right*) reacting with recombinant OspC variants (*y*–axis). The last column (labeled with “-”) is the negative control, showing reactions of sera from un-immunized mice. (*Middle*) Corresponding heat maps. The binding intensity values on the immunoblot images were captured by ImageJ [72]. Values were then normalized by subtracting intensities from the negative controls and by scaling to *z*–scores. (*Bottom*) ARC curves. Some of the most (top-most) and the least (bottom-most) reactive recombinant OspC variants were labeled.

To conclude, testing based on ELISA and immunoblot and using OspC variant-specific mice sera from two species of mice showed the strongest reactions of OspC variants with homologous sera. Reactions of OspC variants with heterologous sera, however, were weaker and inconsistent between experiments and between the two mice species.

### Centroids reacted broadly with naturally infected human and mouse sera

We designed six evolutionary analogs (Supporting Information S1 Alignment) expected to show broad antigenic cross-reactivity with the 16 natural OspC variants using three evolution-based algorithms, including the root algorithm, the consensus algorithm and the centroid algorithm (Fig 4, diagram). Computational analysis of these evolutionary analogs showed their central positions among the OspC sequence diversity (Fig 4, phylogenetic tree). Sequence differences of the evolutionary centroids with the 16 natural variants were more uniform while the consensus analog showed a lower average difference (Fig 4, line plots). The root analog showed the highest average sequence difference as well as the highest variability in sequence differences with the natural variants

**Fig 4.**
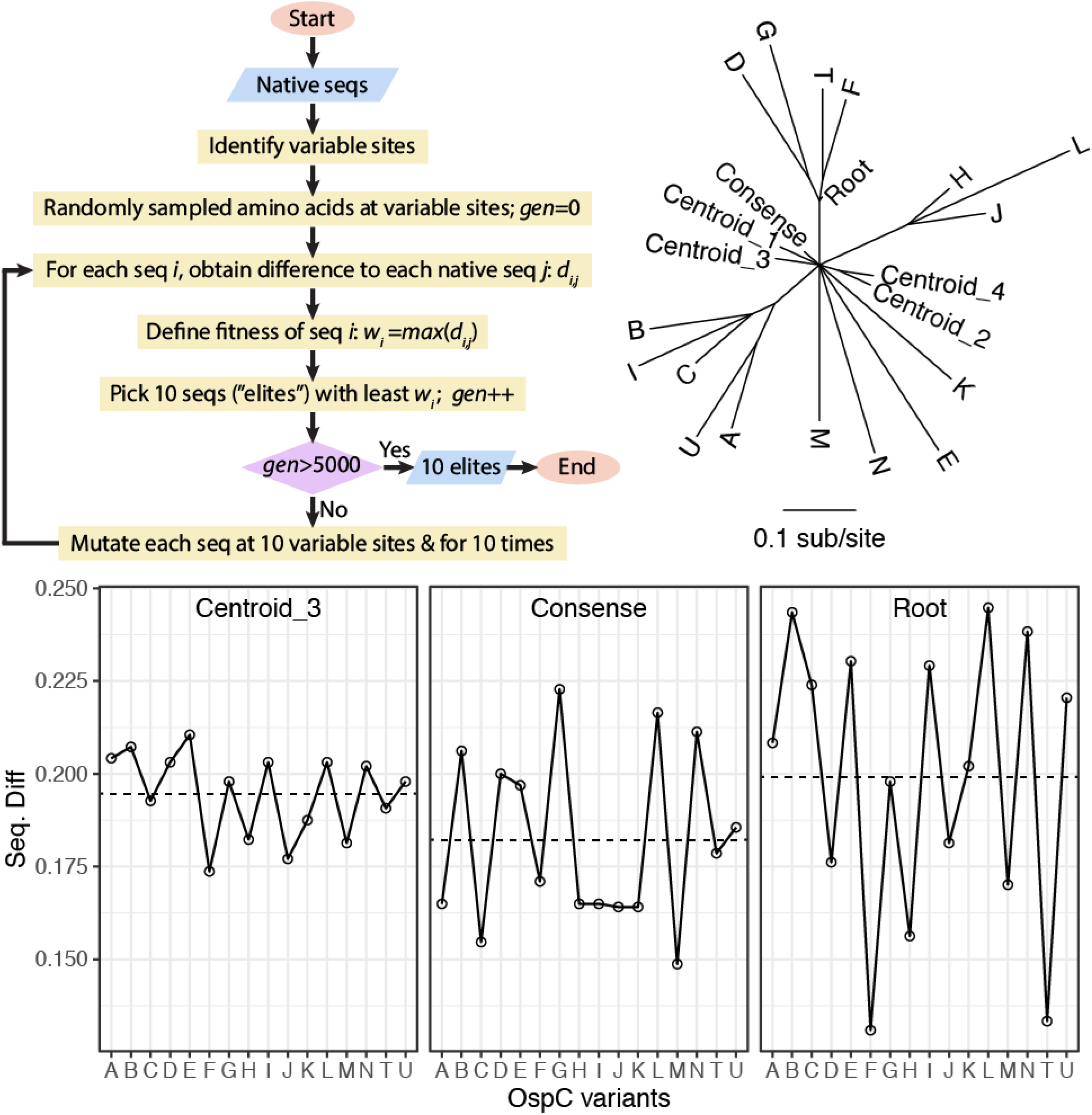
Evolutionary designs of broadly reactive antigens. (*Top left*) The centroid algorithm, which uses genetic algorithm to search for protein sequences with minimal differences to all natural OspC variants (see Material & Methods for details). (*Top right*) A maximum likelihood tree of 16 natural OspC variants (“A” – “U”) common in the Northeast USA and six evolutionarily designed OspC analogs, including the mid-point root sequence (“Root”), a consensus sequence (“Consense”), and four centroids (“Centroid”). All branches are supported by a bootstrap value of 0.8 or above. (*Bottom*) Sequence differences (*y*-axis, in fraction) of three evolutionarily designed OspC analogs with natural OspC variants (*x*-axis).

We cloned, over-expressed, and purified the six evolutionary analogs and the 16 natural variants as recombinant proteins (Supporting Information S2 Fig). Antigenicity of each rOspC was quantified by its reactions with OspC-positive sera (Table 1) from naturally infected human patients (*n*=41) and *P. leucopus* mice (*n*=10) using ELISA (Supporting Information S3d Data; Fig 5). Natural OspC variants (gray bars) reacted with the serum samples with visible variability, so did the root (orange bars) and the consensus (blue bars) analogs. One centroid (“CT1”) reacted poorly with the majority of mice sera and was not further tested with the human sera. In contrast, the other three centroids (“CT2”, “CT3”, and “CT4”) reacted consistently at high levels with all sera. The mouse sera reacted with rOspCs in a more variant-specific manner than the human sera. For example, the mouse sera P03, P04, P06, P08, and P09 reacted strongly with one to four natural rOspC variants while weakly with other natural variants. Although the natural rOspC variants reacted strongly with some of the murine sera, the three centroids reacted consistently high with all murine sera.

**Table 1.** Serum samples

| Label | Host | Source | Description | STTT Interpretation<sup>b</sup> |  |  | C6<br>ELISA<sup>c</sup> |
| --- | --- | --- | --- | --- | --- | --- | --- |
|  |  |  |  | EIA | IgM 23kD<br>Band | IgG 23kD<br>Band |  |
| S01 | Human | CDC | EM<sup>a</sup> convalescence | + | + | + | NA<sup>d</sup> |
| S03 | Human | CDC | EM convalescent | + | + | + | NA |
| S04 | Human | CDC | Non-endemic control | - | + | - | NA |
| S05 | Human | CDC | Non-endemic control | - | - | - | NA |
| S10 | Human | CDC | Neurological Lyme | + | + | + | NA |
| S11 | Human | CDC | EM convalescent | + | + | + | NA |
| S14 | Human | CDC | Fibromyalgia (control) | - | - | - | NA |
| S16 | Human | CDC | Severe periodontitis (control) | - | - | - | NA |
| S18 | Human | CDC | EM convalescent | + | + | + | NA |
| S21 | Human | CDC | Endemic control | - | + | - | NA |
| S22 | Human | CDC | Neurological Lyme | + | + | + | NA |
| S30 | Human | CDC | EM acute | - | + | - | NA |
| T01 | Human | CDC | EM | + | + | + | NA |
| T03 | Human | CDC | Lyme arthritis | + | + | + | NA |
| T04 | Human | CDC | EM | Equ<sup>e</sup> | - | - | NA |
| T05 | Human | CDC | Lyme arthritis | + | + | + | NA |
| T06 | Human | CDC | EM | + | + | + | NA |
| T07 | Human | CDC | EM | Equ | + | - | NA |
| T08 | Human | CDC | EM | + | + | + | NA |
| T09 | Human | CDC | EM | - | - | + | NA |
| T10 | Human | CDC | EM | - | - | - | NA |
| T11 | Human | CDC | EM | + | + | + | NA |
| T12 | Human | CDC | Lyme arthritis | + | - | + | NA |
| T13 | Human | CDC | Neurological Lyme | + | + | + | NA |
| T14 | Human | CDC | EM | - | + | + | NA |
| T15 | Human | CDC | Lyme arthritis | + | + | + | NA |
| T16 | Human | CDC | EM | + | + | + | NA |
| T17 | Human | CDC | EM | + | + | - | NA |
| T18 | Human | CDC | Neurological Lyme | + | + | + | NA |
| T19 | Human | CDC | Cardiac Lyme | + | + | + | NA |
| T20 | Human | CDC | EM | + | + | + | NA |
| T21 | Human | CDC | Neurological Lyme | - | + | + | NA |
| T22 | Human | CDC | EM | + | + | - | NA |
| T23 | Human | CDC | EM | - | - | + | NA |
| T24 | Human | CDC | EM | + | + | + | NA |
| T25 | Human | CDC | EM | + | + | + | NA |
| T26 | Human | CDC | EM | - | + | - | NA |
| T27 | Human | CDC | EM | + | - | + | NA |
| T29 | Human | CDC | EM | + | + | + | NA |
| T30 | Human | CDC | Neurological Lyme | + | + | + | NA |
| T31 | Human | CDC | EM | - | - | - | NA |
| T32 | Human | CDC | Cardiac Lyme | + | + | + | NA |
| H01 | Human | Stony Brook | Late Lyme | NA | NA | NA | 0.924 |
| H04 | Human | Stony Brook | Late Lyme | NA | NA | NA | 1.947 |
| H07 | Human | Stony Brook | Late Lyme | NA | NA | NA | 0.555 |
| H08 | Human | Stony Brook | Late Lyme | NA | NA | NA | 0.260 |
| H09 | Human | Stony Brook | Late Lyme | NA | NA | NA | 0.013 |
| H10 | Human | Stony Brook | Late Lyme | NA | NA | NA | 0.491 |
| P01 | *P. leucopus* | Millbrook, NY | Field reservoir | NA | NA | NA | 0.592 |
| P02 | *P. leucopus* | Millbrook, NY | Field reservoir | NA | NA | NA | 0.266 |
| P03 | *P. leucopus* | Millbrook, NY | Field reservoir | NA | NA | NA | 3.480 |
| P04 | *P. leucopus* | Millbrook, NY | Field reservoir | NA | NA | NA | 0.749 |
| P05 | *P. leucopus* | Millbrook, NY | Field reservoir | NA | NA | NA | 0.910 |
| P06 | *P. leucopus* | Millbrook, NY | Field reservoir | NA | NA | NA | 0.501 |
| P07 | *P. leucopus* | Millbrook, NY | Field reservoir | NA | NA | NA | 0.256 |
| P08 | *P. leucopus* | Millbrook, NY | Field reservoir | NA | NA | NA | 3.046 |
| P09 | *P. leucopus* | Millbrook, NY | Field reservoir | NA | NA | NA | 0.518 |
| P10 | *P. leucopus* | Millbrook, NY | Field reservoir | NA | NA | NA | 0.286 |

<sup>a</sup>EM: erythema migrans, an early-stage Lyme disease
<sup>b</sup>STTT Interpretation: performed and provided by CDC. EIA interpretation based on the VIDAS Lyme IgM and IgG polyvalent assay by bioMérieux, Inc, with the following cutoff values: negative < 0.75, equivocal > 0.75 to < 1.00, and positive > 1.00. IgM and IgG immunoblotting assays by MarDx Diagnostics, Inc.
<sup>c</sup>C6 ELISA: OD450 readings from ELISA using the C6 peptide
<sup>d</sup>NA: not available
<sup>e</sup>Equ: equivocal

**Fig 5.**
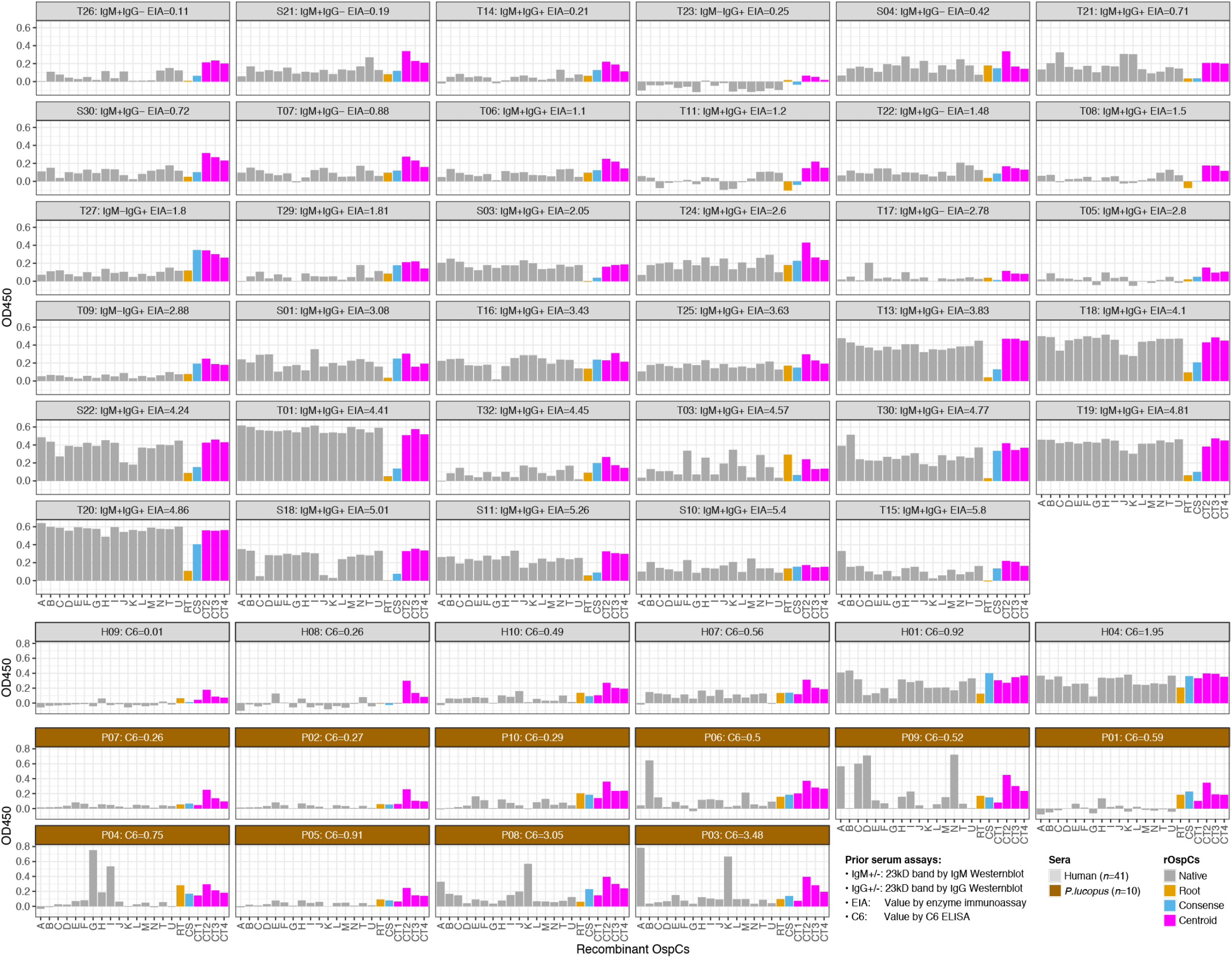
ELISA readings of naturally infected human and mouse sera. Each panel shows OD450 readings (*y*-axis) of a serum sample (listed Table 1) against a panel of rOspCs (*x–*axis). The recombinant OspCs included the 16 natural variants (“A” through “U”), the root sequence (“RT”), the consensus sequence (“CS”), and three centroid sequences (“CT2”, “CT3”, and “CT4”). In the top six rows, sera provided by CDC (*n=*35) were chosen for the presence of the 23 kD band (OspC) in either IgG, IgM, or both types of Westernblot assays. The sera were ordered according to increasing values of the enzyme immunoassay (EIA) readings. The EIA and Westernblot results were provided by CDC [52]. In the bottom three rows, six human sera and ten *P. leucopus* sera were ordered according to increasing values of C6 ELISA results. The recombinant OspCs included an additional centroid sequence (“CT1”).

The antigenic breadths of the OspC variants were further quantified with the use of heat map and ARC curves (Fig 6). In the heat map, the OD450 readings were scaled with respect to individual sera and, subsequently, both the sera (in columns) and the rOspCs (in rows) were grouped according to pair-wise similarities in reactivity (Fig 6, heat map). The three centroids (CT2, CT3, and CT4) showed as a distinct cluster that reacted with the human and mouse sera at levels that were consistently above the average. The ARC curves showed the strong reactions of the three evolutionary centroids with the sera (the top three curves), the weak reaction of the root analog (“RT”, lowest curve), and an average level of reaction of the consensus analog (“CS”, near the zero line). The ARC curve rankings of natural OspC variants (gray lines), indicating relative cross-reactivity or specificity with the naturally infected sera samples, were once more inconsistent with their rankings based on binding with sera from immunized mice (Figs 2 & 3, ARC curves).

**Fig 6.**
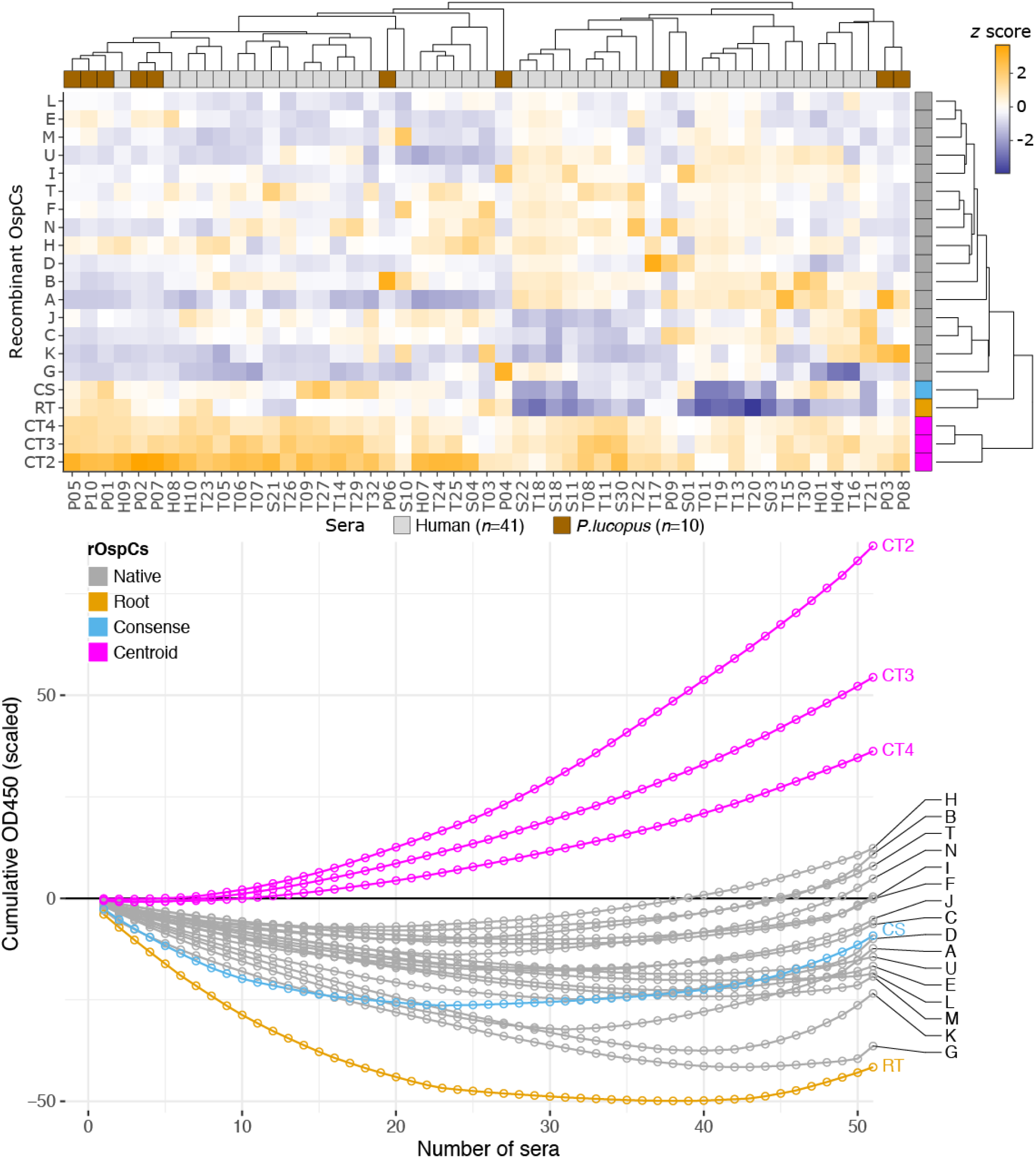
Evolutionary centroids react broadly with OspC-positive sera. (*Top*) A heat map representation of ELISA results of rOspC reactivity with naturally infected sera. Colors represent scaled OD450 values (*z–*scores). (*Bottom*) Antigen reaction characteristics (ARC) curves showing consistently high reactivity of three centroids (CT2, CT3, and CT4), suggesting their broad antigenicity against *B. burgdorferi* strains expressing diverse OspC variants.

We conclude from the ELISA testing that the evolutionary centroids reacted strongly and consistently with naturally infected human and mouse sera, which contained antibodies against a diverse set of antigenically distinct natural OspC variants.

## Discussion

### Natural OspC variants are antigenically maximally diversified

In previous field-based studies, we and others have established an over-abundance of ticks infected by a mixture of Lyme pathogen trains identified by their *ospC* alleles [31,34]. In the present study, we further tested immunological distinctness of diverse *B. burgdorferi* strains coexisting in the Northeast US using field-collected *I. scapularis* ticks. Composition of *B. burgdorferi* strains in individual infected ticks especially in nymphs – having fed on a single blood meal since molting from larvae – faithfully reflects the spirochete composition in reservoir hosts [31,42]. As such, we expected that the frequency of mixed infection by a pair of strains to be lower than expected by chance if cross immunity of reservoir hosts against super-infection by multiple strains was common in nature. The present statistical analysis of coinfection rates reaffirmed an over-abundance of pairs of *B. burgdorferi* strains carrying distinct OspC variants (Fig 1). In conclusion, reservoir hosts of Lyme pathogens tend to be infected by multiple strains, indicating a lack of cross-protective immunity in reservoir hosts. By extension, we conclude the immunological distinctness of *Borrelia* strains carrying different *ospC* alleles in nature.

Experimentation in the lab using *B. afzelii*, a Lyme pathogen common in Europe and Asia, showed that mice immunized with one recombinant OspC variant protected the host from infection by a strain carrying the homologous OspC variant but not by the strain carrying a heterologous OspC variant [24]. These strains, however, differed in genomic background besides the *ospC* locus. Immunological mechanisms by which the host serum neutralizes spirochetes carrying a homologous but not a heterologous *ospC* allele was elucidated using genetic manipulations and immuno-deficient mice, firmly establishing the causal role of OspC molecules in eliciting strain-specific protective humoral immunity in *B. burgdorferi* hosts [25].

By immunizing the C3H mice and the reservoir species *P. leucopus* with recombinant OspC proteins and quantifying antigenic reactions using ELISA and immunoblots, we demonstrated the high antigenic specificities of the full set of natural OspC variants present in a natural *B. burgdorferi* population, as well as a lack of strong and consistent cross-reactivity between heterologous OspC variants (Figs 2 & 3).

To summarize these field-based and lab-based studies, we use the term maximum antigenic diversification (MAD) to describe the immunological distinction of natural OspC variants and their dominant role in maintaining *B. burgdorferi* diversity in nature. Evidence of antigenic separation among natural OspC variants emerged first from population genetic surveys of *ospC* sequence variability and allele frequencies in natural *B. burgdorferi* populations, which showed strong balancing selection driving genetic diversity at the *ospC* locus mediated by ecological mechanisms including immune escape, host-species specialization, or both [26,29,32]. Subsequent whole-genome sequencing revealed frequent recombination among coexisting strains and *ospC* being a recombination hotspot as well as the most polymorphic single-copy gene in the *B. burgdorferi* genome [27,43]. We showed by forward-evolution simulation that the combined forces of homologous recombination and negative-frequency dependent selection were sufficient to explain the seemly paradoxical pattern of the high recombination rate at *ospC* and the sequence hypervariability at the same locus [27].

An epidemiological model offers a more intuitive understanding of the paradox of sustained linkage disequilibrium in the presence of genetic recombination at an antigen locus [3]. Using a token antigen consisting of two bi-allelic epitope sites (e.g., A1 and A2 at site A, B1 and B2 at site B), Gupta *et al* (1996) predicted complete linkage disequilibrium resulting in a population consisting of only A1B1 and A2B2 haplotypes without the crossover A1B2 and A2B1 haplotypes, if it could be assumed that the host antibodies neutralize A1 and A2 (as well as B1 and B2) specifically without cross-reactivity (i.e., anti-A1 not binding A2 and vise versa). This is because, in such a system the A1B1-genotyped microbes would survive the host producing antibodies against A2 and B2, the A2B2-genotyped microbes would survive the host producing antibodies against A1 and B1, but the A1B2− or A2B1-genotyped microbes not survive either host. This simple epidemiological model thus predicts maximum antigenic divergence (two bits of difference between A1B1 and A2B2) when the host immunity is highly allele-specific. This model, known as the strain theory, has been further refined and used to understand the stable coexistence and the temporal persistence of diverse strains in natural pathogen populations including the influenza A (H3N2) virus and malaria [6,44].

Maximum antigenic diversification among natural OspC variants could be understood as a generalized case of the strain theory consisting of ~100 hypervariable epitope sites on the OspC protein sequence. Under this model, the antigenic space defined by the natural OspC variants in *B. burgdorferi* reflects the immunological niches permissible by the reservoir hosts. The large gaps in both the sequence and antigenic spaces among these natural variants are unfilled in natural *B. burgdorferi* populations because spirochetes carrying any recombinant antigens would be vulnerable to neutralization by host immunity. The total number of natural OspC variants in a local *Borrelia* population, however, also reflects the effective pathogen population size, as we have shown with computer simulations [14].

### Evolutionary centroids react broadly with naturally infected sera and specifically with anti-OspC antibodies

We thus designed evolutionary analogs to fill the large genetic and antigenic gaps created as a result of diversifying OspC variants in nature (Fig 4). Among the evolutionary analogs, three centroids designed with a genetic algorithm showed broad reactivity with naturally infected human and mouse sera (Figs 5 & 6). The root and consensus analogs did not show expected broad antigenicity despite their basal tree positions. The centroid algorithm, in contrast, was highly successful and generated three of the four centroid analogs with broad antigenicity. It could be that the antigenic breadth of evolutionary analogs is sensitive to the variance in sequence distances to the natural variants. Indeed, the root analog varied the most in sequence similarity to the natural variants followed by the consensus analog, while the most rigorously computed centroids were the most uniform in their sequence differences with natural variants (Fig 4). However, the centroid algorithm did not succeed uniformly. One of the centroids (“CT1”) showed relatively narrow antigenicity compared to the other three centroids despite being similar in the distribution of sequence distances. To further validate and quantify antigenic breadths of evolutionary analogs, we have begun testing using OspC variant-specific sera from immunized mice.

We performed a preliminary structural analysis by building three-dimensional models of the six evolutionary analogs using a solved OspC structure as the template [45] (Supporting Information S5 Fig). All evolutionary analogs showed high structural similarity with the native template molecule. Validation of structural similarity of evolutionary analogs to native OspC variants requires experimental interrogation using e.g., circular dichroism (CD) and nuclear magnetic resonance spectroscopy (NMR) [46].

We consider the possibility that the consistently high reactivity of the three evolutionary centroids with human and mouse sera was not due to binding with anti-OspC antibodies but with non-OspC antibodies present in the sera. To rule out the possibility of non-OspC specific reactions, we compared the reactivity of evolutionary analogs with the reactivity of non-OspC antigens as negative controls. Specifically, we performed regression analyses of the reactions of the five evolutionary analogs and non-OspC antigens with the average levels of anti-OspC antibodies present in the sera (Fig 7). We used the average OD450 readings of a serum against the 16 natural variants as a proxy of the total amount of anti-OspC antibodies present in the serum. The non-OspC antigens, e.g., BSA and the C6 peptide (a part of the VlsE protein), were not expected to react strongly or consistently with anti-OspC antibodies.

**Fig 7.**
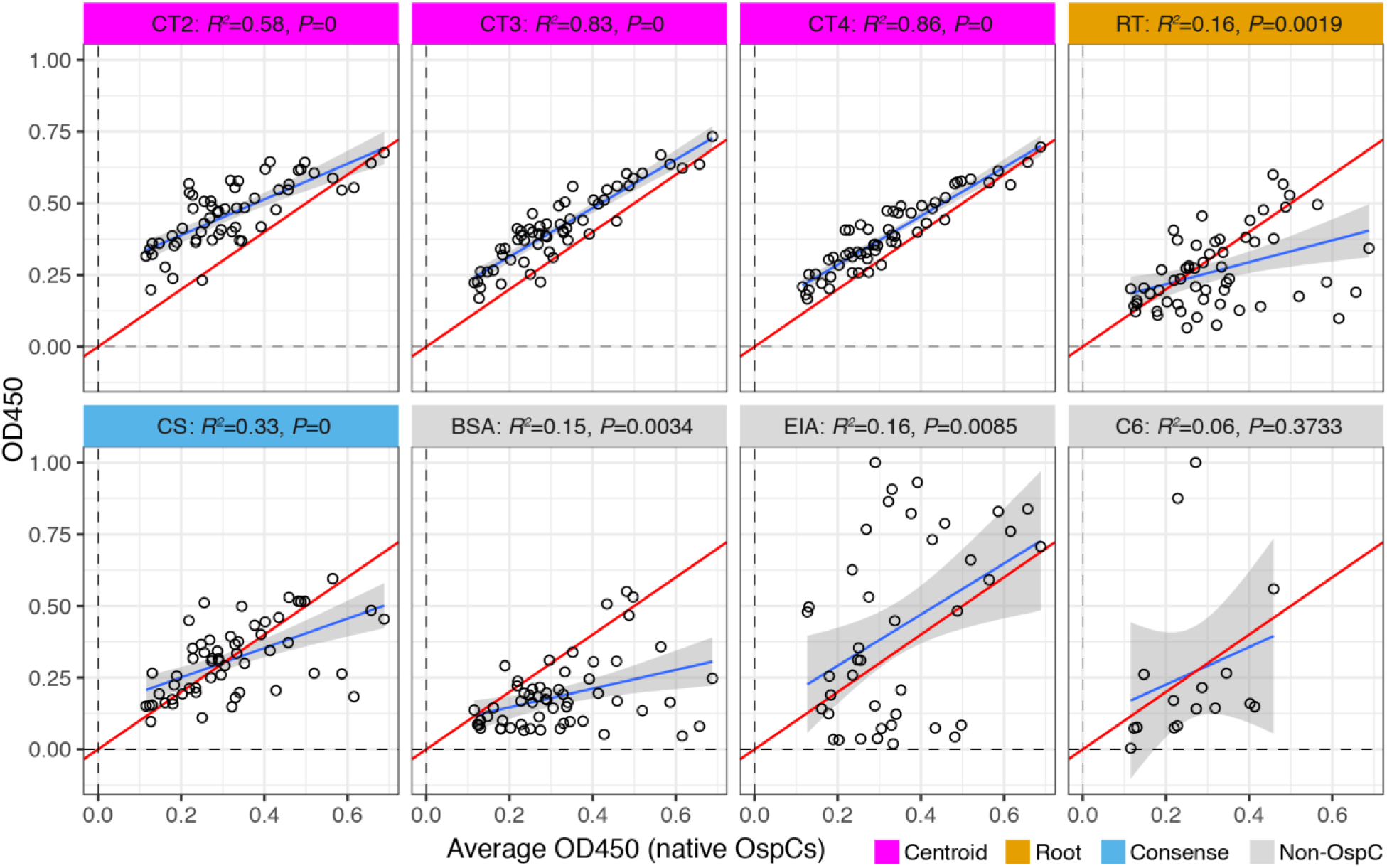
Evolutionary centroids react specifically with anti-OspC antibodies. Each panel represents a specific antigen (listed in heading). Within a panel, points represent individual sera. The *x*-axis shows the average OD450 reading of a serum against the panel of 16 natural rOspC variants, as a proxy for the total amount of anti-OspC antibodies present in sera. The *y*-axis shows the OD450 readings of an antigen’s reaction with the sera. Thus, a regression line (in blue) represents the degree by which an antigen’s reaction with the sera is attributed to its reaction with anti-OspC antibodies. The strength of correlation is measured by *R^2^* (i.e., proportion of explained variance) and the statistical significance by the *P* values (both shown in the panel heading). The red line indicates the 1:1 ratio (i.e., equivalence) between the two readings. The panels show reactions of five evolutionary analogs (“CT2”, “CT3”, “CT4”, “RT”, and “CS”) and an non-OspC negative control (“BSA”, for bovine serum albumin) with *n*=57 sera (including OspC-negative sera, Table 1). Two additional non-OspC negative control antigens included the whole-cell lysate used in EIA testing (“EIA”) for reaction with *n*=35 sera provided by CDC, and the VlsE C6 peptide (“C6”) used in ELISA for reaction with *n*=16 sera. The EIA and C6 ELISA values (Table 1) were re-scaled to values between zero and one. Reactions of the three centroids were strongly associated with the levels of anti-OspC antibodies (high *R^2^* and low *P* values), while all negative controls showed weak correlation, a lack of statistical significance, or both.

As expected, only a small portion of the reactivity of non-OspC antigens may be attributed to their binding with OspC-specific antibodies (~15% for BSA and not significant for C6). The root and consensus analogs also reacted poorly (although significantly) with anti-OspC antibodies, with 16% and 33% explained variance respectively. In contrast, the three centroids showed 58-86% explained variance with high statistical significance. We conclude that although binding with non-OspC antibodies may have occurred in ELISA testing of the evolutionary analogs, such non-specific binding was not a major contributor to the high and broad reactivity of the evolutionary centroids. In other words, the evolutionary centroids were not only broadly cross-reactive with natural OspC variants but also highly specific in their recognition of anti-OspC antibodies present in human and mouse sera.

The regression analysis revealed an upward bias of OD450 values for CT2, CT3, and CT4 centroids (Fig 8). Indeed, evolutionary centroids showed OD450 values higher than those of any natural variant in ELISA tests of OspC-negative sera and weakly reactive sera (e.g., sera with low EIA or C6 ELISA values, Fig 5, bar plot). It remains to be investigated whether the bias is a feature of evolutionary centroids or due to experimental biases. Nevertheless, the centroid antigens displayed consistently robust reactions even in cases where their binding levels were only about half of the levels of the most reactive natural OspC variants (see reaction profiles of the mouse sera P03, P04, P06, P08, and P09 in Fig 5, bar plot). It is also not clear why human sera tended to show weaker variant-specific reactions compared with the *P. lucopus* sera (Fig 4, bar plot). Humans are accidental host of *B. burgdorferi*. It is conceivable that the spirochete cells are not well adapted to the human host environment and tend to disintegrate upon entry. As a result of such mal-adaptation to the human host, conserved domains of *B. burgdorferi* surface antigens (e.g., the C6 domain of the VlsE protein) may elicit antibody responses in humans but not in natural reservoir species [47]. Further testing using, e.g., more replications, more precise antigen load control, and sera from mice immunized with natural OspC variants, is necessary for improved quantification of the antigenic breadth of the evolutionary centroids.

**Fig 8.**
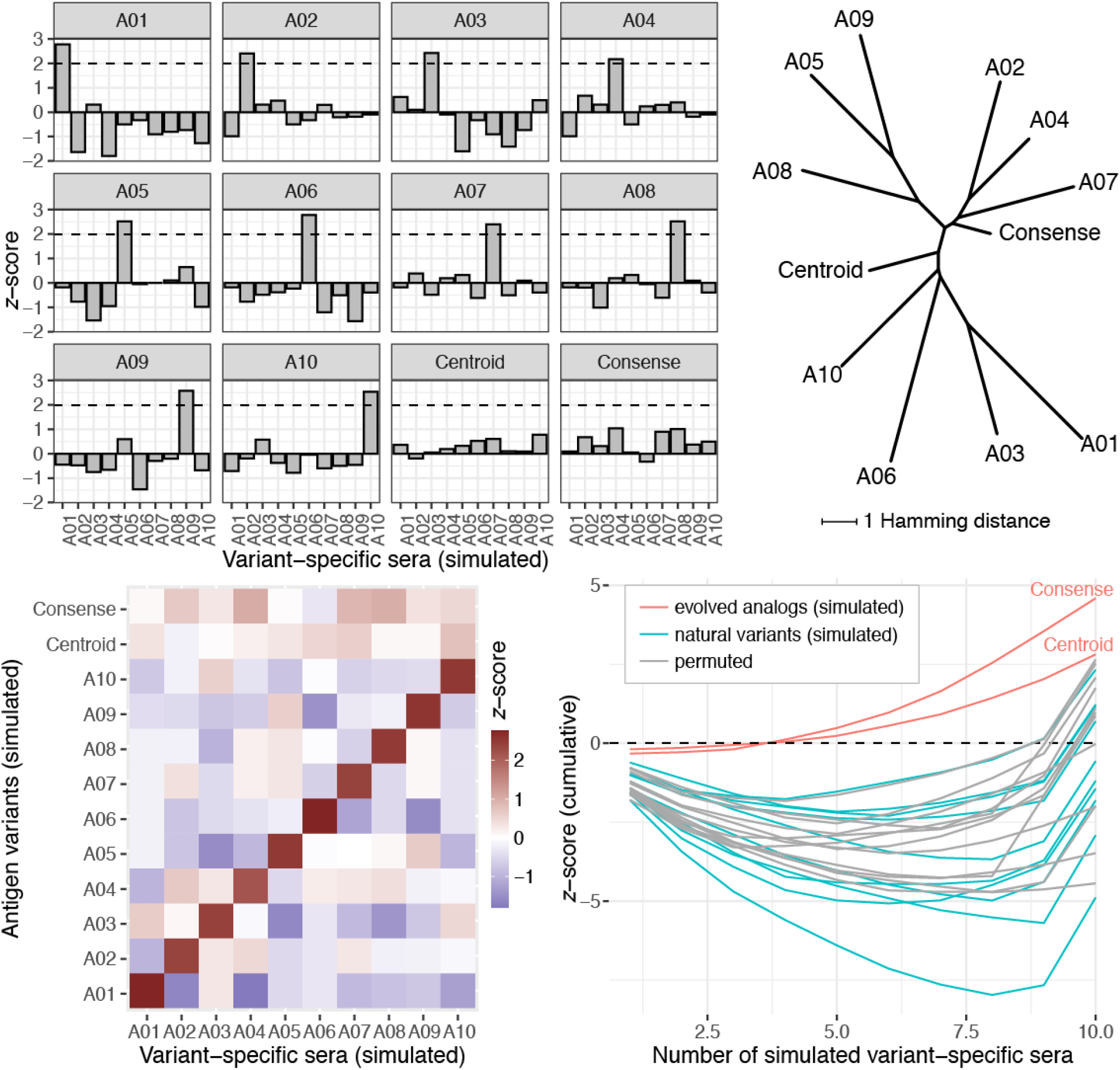
Simulated divergent antigens and evolutionary analogs. The simulated population initially contained ten antigen variants, each represented by a 20-bit long randomly generated binary string. Each bit represented a variable antigen site. Using a genetic algorithm (Supporting information S4 R Markdown), we created a simulated population consisting of alleles “A01” through “A10” with maximally divergent sequences (Table 2). Subsequently, a consensus sequence (“Consense”) was created from majority bits at individual positions. The centroid sequence (“Centroid”) was generated by minimizing maximum sequence differences to the natural variants using a second round of genetic algorithm. Antigenic reactivity of a simulated variant *i* to a simulated variant *j*-specific serum was assumed to be proportional to the sequence similarity between the two variants. Levels of antigenic activity were normalized to *z*-scores with respect to each simulated variant-specific serum. (*Top left*) Each panel shows levels of reactivity (*x–*axis) of a simulated variant to simulated variant-specific sera (*y–*axis). (*Top right*) A neighbor-joining tree of simulated variants based on pair-wise Hamming distances. (*Bottom left*) A heat map representation of the levels of antigenic activity between the simulated variants (*y–*axis) and the simulated variant-specific sera (*x–*axis). (*Bottom right*) Each line shows cumulative *z–*scores (*y–*axis) of a simulated natural variant (cyan), an evolutionary analog (red), or a natural variant after one round of permutation of *z*-scores among the simulated natural variants (gray).

### Mechanism of broad antigenicity of evolutionary centroids: a simulation model

To explore immunological and molecular mechanisms underlying the broad antigenicity of evolutionary centroids, we computationally simulated maximum antigen diversification (MAD) and evolutionary centroids using genetic algorithms (Supporting Information S4 R Markdown; Material & Methods). Simulated maximally diverged antigens (Table 2) were validated by constructing a neighbor-joining tree based on pair-wise Hamming distances, which showed the central positions of the consensus and centroid variants (Fig 8, tree). Simulation results were further validated by tabulating pairwise Hamming distances into a distance matrix, which showed a narrow range of distances (6 to 9) for the centroid, and a wider range of distances (4 to 11) for the consensus, and large distances (6 to 15) between simulated natural variants (Table 2). We obtained *z*-scores by normalizing the sequence similarities with respect to individual simulated natural variants. As such, we were able to compare and show a high resemblance between the simulate results and experimental results using immunized mice (Figs 2, 3 & 8). For example, both the simulated and experiment-derived results showed highly specific bindings (*z* > 2.0) between homologous variants (Figs 2 & 8). Both the simulated and experiment-derived results showed the broad antigenicity of evolutionary analogs with heat map and ARC curves (Figs 6 & 8). Importantly, the simulated MAD population, mirroring experimental results, revealed that the broad antigenicity of evolutionary analogs was a result of consistently above-average (*z* > 0) reactivity even as their cross-reactions with any particular natural antigens remained uniformly lower than the strongest reactions between homologous reactions (*z* < 1) (Fig 8, bar plots).

**Table 2.**
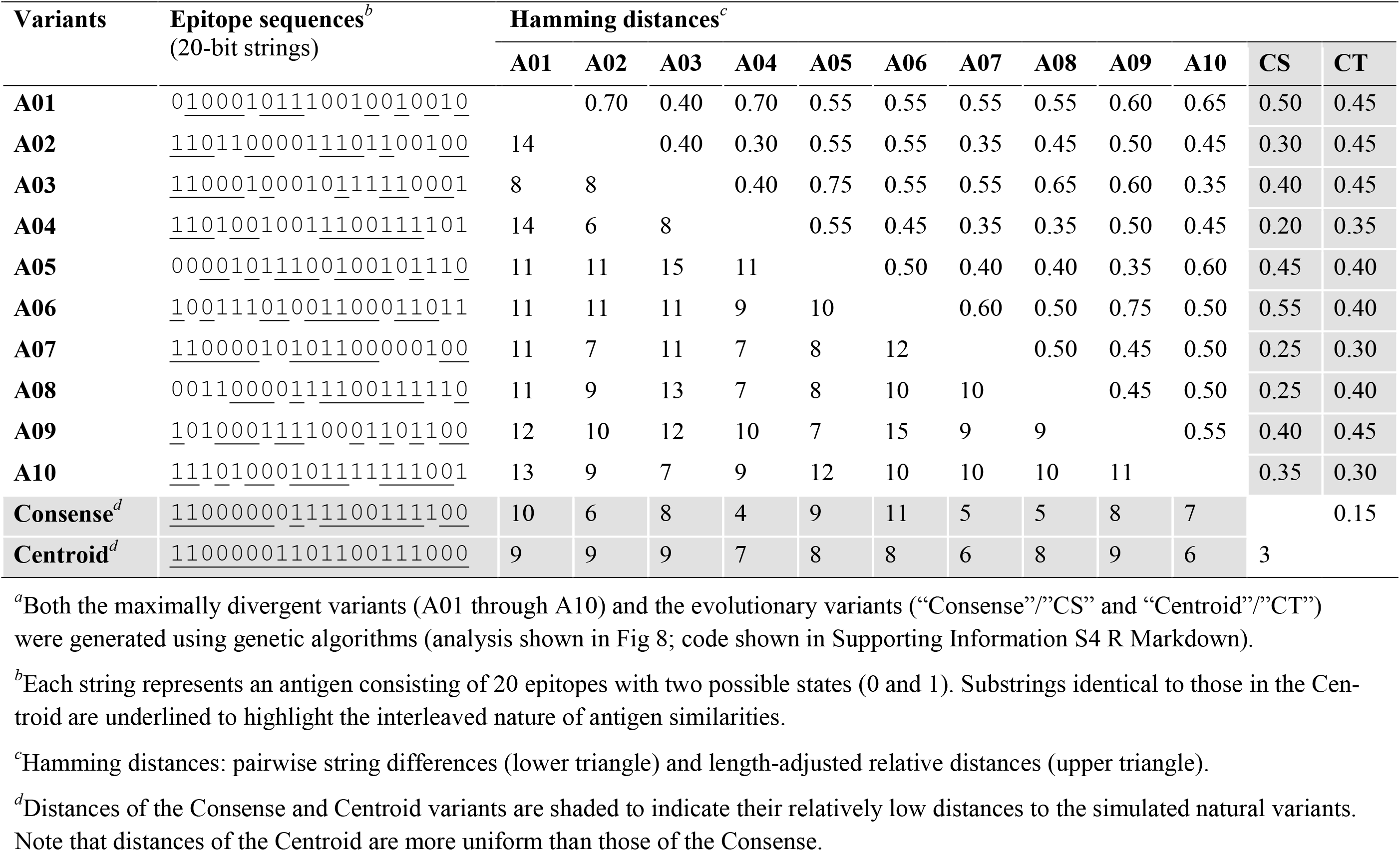
Simulated maximally diverged variants and evolutionary analogs*^a^*

### Implications to diagnostic and vaccine development

A new class of broad-spectrum diagnostics and vaccines could be designed by countering the evolutionary trend of maximum antigenic divergence in local *B. burgdorferi* populations. In diagnosis of Lyme disease, the standard two-tied testing (STTT) is based on EIA and immunoblots and lacks sensitivity for patients who develop acute erythema migrans, an early-stage Lyme disease [48,49]. The newly recommended modified two-tied testing (MTTT) protocol, following a two-step EIA without immunoblot, improved the sensitivity of detecting early Lyme disease cases [50,51]. Previously, we proposed the use of multiple OspC variants for sensitive detection of *B. burgdorferi* infections [37]. The centroid antigens, with their broad reactivity with sera including those with low EIA readings, are novel diagnostic candidates if they pass specificity tests [52].

Currently there is no human-use vaccine against Lyme pathogens in the US [53–55]. Design of OspC-based vaccines focuses on identification of immuno-dominant epitopes in individual OspC variants and concatenating them into linear multivalent super-antigens [35,56,57]. A multivalent vaccine consisting of as many as eight OspC-type specific epitopes has been developed for canine use [38]. Due to the large number of OspC variants co-circulating in a local endemic area (e.g., ~20 in the Northeast US) and uncertainties in epitope identification, it remains unclear the efficacy of chimeric vaccines to elicit broadly protective immunity in humans [39,58].

Experiments are under way to measure the immunogenicity of evolutionary centroids *in vivo* as well as to evaluate their vaccine efficacy upon challenging the immunized mice with infected ticks carrying diverse *B. burgdorferi* strains. If validated and used as reservoir-targeted vaccines [59], the centroid antigens have the potential to reduce spirochete loads in natural reservoir hosts by eliciting immunity against all Lyme pathogen strains.

Vaccines based on centroid antigens would be similar to the COBRA (Computationally Optimized Broadly Reactive Antigen) vaccines against influenza viruses [8]. Both approaches are based on principles of antigen evolution and use automated computational design. While the COBRA vaccines are based on consensus sequences, the centroid-antigen design uses genetic algorithms. The consensus OspC analog failed to display broad antigenicity in the present study. By enforcing minimal sequence differences with all natural variants, the centroid algorithm was the most effective in broadening antigen cross-reactivity (Fig 4).

Stable coexistence of antigen variants like OspC variants in *B. burgdorferi* is widespread in natural pathogen populations. The Dengue viral populations consist of four antigenically distinct serotypes associated with sequence variations of the envelop protein [60]. The influenza B viral populations contain two evolutionary lineages associated with sequence variations of hemagglutinin [61]. The malaria parasite populations are structured into antigenic groups associated with genetic variations of the *var* genes encoding an erythrocyte membrane protein [6]. If these pathogen strains indeed represent ecological niches shaped by host immunity [3,62], evolutionary centroids would be a novel and effective strategy against a broad range of microbial pathogens.

## Materials & Methods

### Co-occurrences of OspC variants in field-collected Ixodes ticks

We tested the immunological distinction of *B. burgdorferi* strains based on their co-occurrences in individual *Ixodes scapularis* ticks. We expected that immunologically distinct strains could infect a single host and thus tended to co-infect a single tick while immunologically similar strains tended to be found in different ticks. Compositions of *B. burgdorferi* strains infecting individual ticks reflect those in the reservoir hosts [42]. The analysis was based on the presence and absence of 20 Lyme pathogen strains infecting *n*=119 *I. scapularis* ticks collected from New York State during 2015 and 2016 [41]. The over- or under-abundance of a pair of strains (*i* and *j*) was quantified as the fold change of the observed over the expected counts: 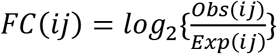. Statistical significance of the relative abundance was obtained from a null distribution generated by permuting the occurrences of a pair of strains among infected ticks 1000 times, keeping the total occurrence constant.

### Evolutionary algorithms for designing broadly reactive synthetic OspC

Protein sequences analogous to natural OspC variants were optimized for broad cross-reactivity using the following evolutionary algorithms. All algorithms aimed to generate sequences close to the center of the diversifying natural OspC variants (Fig 4). First, we reconstruct the hypothetical ancestral sequence at the mid-point root of the phylogeny of natural OspC variants using RAxML [63] (the “Root Algorithm”).

Second, we obtained a consensus sequence consisting of 20% majority residues at aligned sequence positions of natural OspC variants using the consensus method implemented in the Bio::SimpleAlign module of the BioPerl library [64] (the “Consensus Algorithm”).

Third, we used genetic algorithm to generate sequences with minimal distances to natural OspC variants (the “Centroid Algorithm”) (Fig 4). Briefly, we extracted amino acids at variable positions from 16 aligned natural OspC sequences commonly found in Northeast USA [26,31] (Supporting Information S1 Alignment). An initial seed population of random antigen sequences (e.g., *n*=10) were generated by sampling amino acids with uniform probabilities of distinct residues present at each variable site. For each randomly generated sequence *i*, we calculated its differences (*d_ij_*, *j*=”A” through “U”) to the 16 natural variants. We defined the fitness of this sequence as the maximum value among its differences to all 16 natural sequences: *w_i_* = *max*(*d_ij_*). This fitness measured its overall sequence similarity to the natural variants: the lower the *w_i_* the higher its overall similarity to the natural variants. The top ten most similar antigen sequences at each generation were retained and others were discarded. Each elite sequence was then allowed to “reproduce” and generate ten “gametes” with mutations at randomly selected ten variable sites. The above process was repeated (e.g., for 5000 generations) to progressively lower the *w_i_* values, after which the final output included ten elite centroids that were the most similar to all 16 natural OspC variants. The centroid algorithm was implemented using BioPerl library in Perl [64] and the DEAP package in Python [65]. The top four optimized centroid sequences were cloned, over-expressed, and purified for immunological assays of antigenic breadth.

### Gene synthesis, protein over-expression, and protein purification

DNA sequences encoding the natural and synthetic OspC variants were codon-optimized, synthesized, and cloned into the pET24 plasmid vector, which was then used to transform the *Escherichia coli* BL21 cells. All DNA work was performed by a commercial service (GeneImmune Biotechnology Corp., Rockville, MD, USA). We designed the OspC constructs by excluding the first 18 residues encompassing the signal peptide and by adding a 10 × Histidine-tag on the N-terminus. These modifications were necessary for over-expression of OspC proteins in *E.coli* and to facilitate OspC purification [46].

For each *E. coli* strain containing a cloned *ospC* gene, a single colony was selected to inoculate 4 ml Luria-Bertani (LB) broth (Thermo Fisher Scientific, Waltham, MA, USA) containing vector-specific selective antibiotics (25 μg/ml for Kanamycin or 50 μg/ml for Ampicillin). The seeded culture was incubated overnight at 37° with vigorous shaking (250 rpm). A portion of the overnight culture was transferred into 50 ml fresh pre-warmed LB broth containing 0.4% glucose and the selective antibiotics. The culture was incubated at 37° with vigorous shaking until reaching exponential growth indicated by an OD_600_ of approximately 0.8 as measured by the NanoDrop Spectrophotometer (Thermo Fisher Scientific, Waltham, MA, USA). Expression of the cloned *ospC* was induced by adding isopropyl β-d-1-thiogalactopyranoside (IPTG) to a final concentration of 0.25 – 0.5 mM and by incubation overnight at 25°. Cells were collected by refrigerated centrifugation at 4° and 7200 rpm for 15 min, re-suspended in a lysis buffer containing 0.2 mg/ml lysozyme, 20mM Tris-HCl (*p*H 8.0), 250 mM NaCl, and 1mM dithiothreitol (DTT). After incubation for one hour at 4°, cells were further lysed by sonication until the solution become translucent. The lysate was centrifuged in refrigeration at 12,000 rpm for 20 min and the supernatant was withdrawn.

The recombinant proteins were purified using nickel sepharose beads (Ni-NTA, Thermo Fisher Scientific, Waltham, MA, USA). The lysate supernatant from the 50 ml culture was mixed with 300 μl Ni-NTA beads and incubated overnight at 4° in the lysis buffer supplemented with 5 mM imidazole. The lysate-bead mixture was then loaded into a chromatography column and washed with 12 times the bed volume of the lysis buffer containing 25 mM imidazole. The purified protein was eluted with 6 times the bed volume of the lysis buffer containing 500 mM imidazole. The elution was dialyzed to remove imidazole in phosphate-buffered saline (PBS, *p*H7.4) containing 1mM DTT and 20% glycerol.

The amount and purity of recombinant proteins were examined using the sodium dodecyl sulfate polyacrylamide gel electrophoresis (SDS-PAGE) containing 12% gel following the standard protocol. The PageRuler™ Prestained 10 −180 kDa Protein Ladder (Thermo Fisher Scientific, Waltham, MA, USA) was used to mark molecular weights. The gel was stained in 0.08% Coomassie Blue and de-stained in 45% methanol and 10% acetic acid. Concentration of the final purified protein solution was quantified using the Pierce™ Bradford Protein Assay Kit (Thermo Fisher Scientific, Waltham, MA, USA).

### Sera from naturally infected hosts and immunized mice

The majority of human serum samples were provided by the Centers for Disease Control and Prevention (CDC) (Table 1). The human sera originated from patients diagnosed with early to late Lyme disease or from healthy individuals in endemic and non-endemic regions in the USA [52]. The CDC sera panel was previously screened using the standard two-tied diagnostic testing (STTT) for the presence of antibodies against *B. burgdorferi,* including IgM, IgG, or both antibodies against OspC (the 23 kD band) [49]. The CDC sera panel was custom compiled for the present study. Ten serum samples from Lyme disease patients were originally collected by the Stony Brook University Health Science Center, NY, USA. Ten serum samples were obtained from the natural reservoir of *B. burgdorferi*, the white-footed mouse (*Peromyscus leucopus*) from Milbrook, NY, USA. The latter human and mouse sera were screened for exposure to *B. burgdorferi* using the C6 ELSIA (Immunetics, Boston, MA, USA).

Sixteen recombinant OspC were cloned in the pET9c plasmid, the proteins were expressed in *E. coli* BL21 (DE3) pLysS and purified under native conditions by ion exchange chromatography using Q-Sepharose Fast Flow (GE Healthcare, Sweden) as described previously [66]. C3H/HeJ mice (*Mus musculus*) and white-footed mice (*P. leucopus*) were immunized with 10-20 μg of each of the 16 individual purified natural recombinant OspC proteins. Briefly, mice received a dose of recombinant protein on Day 1 and Day 14, and on Day 28 they were euthanized and blood collected by heart puncture. Animal experimentation followed the protocols approved by the Animal Care and Use Committee of University of Tennessee Health Science Center.

### Immunological assays

Immunoblot assays of OspC variant-specific sera were performed using a MiniSlot/MiniBlotter 45 system (Immunetics, Boston, MA, USA). Briefly, a PVDF membrane (Millipore, Billeirca, MS, USA) was mounted on the MiniSlot and 25 μg of each purified protein was loaded individually into its parallel channels. The proteins were immobilized onto the PVDF membrane after the excess solution was removed by vacuum aspiration, resulting in a deposit of horizontal parallel stripes of antigens. The membrane was released from the MiniSlot and blocked in 10% skim milk (Difco, Sparks, MD, USA) for 2 hours at room temperature. After blocking the membrane was rotated by 90 degrees and placed in the MiniBlotter 45. Diluted mouse serum (1:100 to 1:1000 in 3 % of milk in TBS buffer with 0.5% Tween 20, 150 μl) was deposited in the individual vertical lanes of the Miniblotter and was incubated for 1 hour at room temperature. The membrane was washed three times with TBS containing 0.5% Tween 20 and was incubated with goat anti-mouse IgG conjugated with alkaline phosphatase (1:2000) (Kirkegaard & Perry Laboratories [KPL], Gaithersburg, MD, USA) for 1 hour at room temperature. The BCIP/NBT Phosphatase Substrate (KPL) was used to visualize the signal. Serum of non-immunized mice and BSA were used as negative controls.

Sera from naturally infected hosts were individually tested for reactivity with recombinant OspC proteins (rOspCs) using enzyme-linked immunosorbent assay (ELISA). Specifically, a 96-well MICROLON 600 plate (USA Scientific, Inc., Ocala, FL, USA) was loaded in each well with 100 μl PBS containing 10 μg/ml of a rOspC and incubated overnight at 4°. The coated plate was washed three times using PBS containing 0.1% Tween20 (PBS-T buffer) and blocked with 200 μl PBS-T buffer containing 5% milk for one hour at 37°. After washing three times with the PBS-T buffer, 100 μl serum sample diluted in PBS by a factor between 1:100 to 1:1000 was added to each well and incubated for one hour at 37°. After washing three times with the PBS-T buffer, 100 μl diluted horseradish peroxidase (HRP)-conjugated secondary antibodies was added to each well. We used the Goat Anti-Human IgG/IgM (H+L) (Abcam, Cambridge, UK) diluted by a factor of 1:50,000 for assays of human sera and the Goat Anti-*Peromyscus lucopus* IgG (H+L) (SeraCare Life Sciences, MA, USA) diluted by a factor of 1:1,000 for assays of *P. leucopus* sera. After incubation for one hour at 37° and washing with PBS-T buffer, 100 μl TMB ELISA Substrate Solution (Invitrogen™ eBioscience™) was added. The enzyme reaction proceeded for 15 – 30 min at room temperature and was terminated with 1 M sulfuric acid. Binding intensities were measured at the 450 nm wavelength using a Spectra-Max^®^ i3 microplate reader (Molecular Devices, LLC, CA, USA).

### Statistical analysis of OspC cross-reactivity

We tested immunological specificity of natural OspC variants by a re-analysis of the data sets that were the basis of a previously published study [37]. The data sets consisted of ELISA readings of the binding between 16 common recombinant OspC variants segregating in North-east United States *B. burgdorferi* populations with antisera (*n*=15) from C3H/HeJ mice (*Mus musculus*) artificially immunized with individual OspC variants. Sera from uninfected mice were used as negative controls. Additional data sets were ELISA readings of the 16 recombinant OspC variants binding with sera from naturally infected hosts, including the whited-footed mice (*Peromyscus leucopus*, *n=*43), dogs (*Canis lupus*, *n=*25), human Lyme disease patients from the United States (*n=*25), and human patients from Europe (*n=*40).

First, we rescaled the raw ELISA readings into normalized *z*-scores: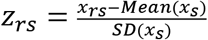, where *x_rs_* is the binding value (OD_450_ reading) of the recombinant OspC *r* with the serum *s* while Mean(*x_s_*) and SD(*x_s_*) represent, respectively, the mean and the standard deviation of OD_450_ readings of all recombinant OspC variants with the serum *s*. The rescaling was employed to account for observed systematic differences among sera samples in non-specific, background bindings introduced by e.g., varying serum dilution factors.

Second, we designed a novel statistical measure to quantify and compare antigenic specificities of OspC variants. The antigenic reaction characteristic (ARC) of an OspC variant is a curve of cumulative binding values (e.g., scaled OD_450_ readings) against the cumulative number of serum samples. A highly specific antigen variant would show a low-lying ARC curve indicating consistently low bindings with sera except those exposed to the variant itself. A broadly cross-reactive antigen variant, in contrast, would show an elevated ARC curve indicating consistently high bindings with sera. As such, the <u>a</u>rea <u>u</u>nder an ARC <u>c</u>urve (AUC) corresponds to a higher cross-reactivity (or lower specificity) of an OspC variant. The ARC curve is similar to the receiver operating characteristic (ROC) curve, which quantifies the specificity of a detection or classification method by plotting the cumulative number of true positives against the cumulative number of false positives [67]. Statistical significance of an ARC curve was evaluated by comparing with curves generated by randomly permuting the scaled OD_450_ readings among all samples.

### Simulation of maximum antigen separation and evolutionary centroids

To explore immunological and molecular mechanisms underlying the broad antigenicity of evolutionary centroids, we computationally simulated maximum antigen diversification (MAD) and evolutionary centroids using genetic algorithms. Following the multiple-epitope extension of the Gupta *et al* (1996) model [3], we represented antigen variants in a pathogen population as 20-bit binary strings (Table 2). We used genetic algorithm to search for a sample of maximally separated antigen variants to represent a MAD population. The searching was performed using the GA package in R [68] and with a fitness function *fit* = max(*d_i,j_*), where *d_i,j_* represents pair-wise Hamming distances between simulated variants. A MAD population is created when this fitness function is maximized. To search for centroid variants, a separate genetic algorithm was used with a fitness function: *fit* = max(*d*_*i,j*=1:10_), where *i* is an artificial variant and *j* is one of the ten simulated natural variants. A centroid allele is found when this fitness function is minimized. Evolutionary analogs were validated with a neighbor-joining tree based on pair-wise Hamming distances. An R markdown of the simulation protocol is included as Supporting Information S4.

We note the similarity between the simulated maximally diversified antigens and the well-separated binary codewords in a self-correcting Hamming code [69,70]. We further note that the problem of finding centroids given a set of strings is known as the Closest String or Hamming Centroid problem in computer science [71]. While the Hamming code and the Hamming centroid problems could be solved with exact algorithms, we solved both problems stochastically with genetic algorithms in the present study.

### Data and code availability

Datasets are provided in Supporting Information S3 Data. Source codes are available in a Github repository (https://github.com/weigangq/ag-div).

## Authors contributions

Conceptualization: Maria Gomes-Solecki, Weigang Qiu

Model development: Saymon Akther, Brian Sulkow, Weigang Qiu

Experimental Investigations: Lia Di, Saymon Akther, Larisa Ivanova, Bing Wu

Software implementation: Lia Di, Edgaras Bezrucenkovas

Data analysis: Lia Di, Edgaras Bezrucenkovas, Brian Sulkow, Larisa Ivanova, Weigang Qiu

Data curation: Edgaras Bezrucenkovas, Weigang Qiu

Funding acquisition: Maria Gomes-Solecki, Weigang Qiu

Supervision: Maria Gomes-Solecki, Weigang Qiu Writing – original draft: Weigang Qiu

Writing – review and editing: Lia Di, Saymon Akther, Brian Sulkow, Edgaras Bezrucenkovas, Maria Gomes-Solecki

## Acknowledgements

This work was supported by the Public Health Service awards AI139782 (to WGQ) and AI072810, AI074092 and AI155211 (to MGS) from the National Institute of Allergy and Infectious Diseases (NIAID) of the National Institutes of Health (NIH) of the United States of America. Additional funding includes the grant CK000107 (to MGS) from the US Centers for Disease Control and Prevention (CDC). The content of this manuscript is solely the responsibility of the authors and does not necessarily represent the official views of NIAID, NIH, or CDC. Saymon Akther is supported in part by the Doctoral Program in Biology of the City University of New York. We thank Dr Mirella Salvatore (Weil Cornell Medical College) for introducing us to the study of influenza B viruses. Dr Christopher Sexton and Dr Jeannine Petersen (Division of Vector-Borne Diseases, CDC) prepared a custom panel of human sera for this study. Roman Shimonov and Justin Hiraldo (both of Hunter College) digitalized immunoblot images.

## Supporting information caption

**S1 Alignment. Aligned sequences of natural OspC variants and evolutionary analogs**

The alignment includes sequences of 16 natural OspC variants (“A” through “U”) prevalent in the Northeast US [26,31] and the six evolutionary analogs (“Root”, “Consense”, and “Centroids”). Signal peptide (N-terminal 18 residues) was removed from each recombinant protein construct. The constructs also contained a 10X histidine tag at the N-terminus (not shown).

**S2 Figure. SDS-PAGE images of purified recombinant OspC proteins**

(*Top*) Purified 16 recombinant natural OspC variants used in ELISA and immunoblot assays with sera from immunized C3H and *P. leucopus* mice. (*Bottom*) Purified 16 recombinant natural OspC variants (“A” through “U”) and six evolutionary analogs used in ELISA including four centroids (“CT1” through “CT4”), one phylogenetically reconstructed root sequence (“RT”), and one consensus sequence (“CS”). Purified rOspCs show as bands near the 25 kD molecular weight marker. Purified OspC molecules tend to dimerize due to presence of two cysteine residues, producing fainter bands between the 40 kD and 55 kD markers.

**S3 Data.**

These are Excel sheets including the following five datasets: (a) counts of presence and absence of OspC variants in 119 infected ticks collected from four study sites in New York, USA [31], used for plotting Fig 1; (b) mean OD450 readings (averaged from two replicates) from ELISA of 16 recombinant OspC proteins with 15 variant-specific sera [37], used for plotting Fig 2; (c) digitalized binding intensity values of two immunoblots of 16 recombinant OspC proteins with 16 variant-specific sera from C3H and *P. leucopus* mice, used for plotting Fig 3; (4) OD450 readings from ELISA of 16 natural rOspCs and six evolutionary analogs with OspC-positive sera from human patients (*n*=41) and *P. leucopus* mice (*n*=10), used for plotting Figs 5 & 6; (5) each row includes OD450 reading of a variant with an serum, long with the average value of OD450 readings of the serum with the 16 natural OspC variants, used for plotting Fig 7.

**S4 R Markdown. Simulation of maximal antigen divergence and evolutionary analogs using genetic algorithms**

Part 1: Generate maximally divergent 20-bits long binary sequences using genetic algorithm implemented in the GA package [68]. Part 2: Find broadly cross-reactive centroids, again using genetic algorithm implemented in the GA package [68]. Part 3: Print simulation results and validate with a neighbor-joining tree.

**S5 Figure. Structural superimposition of evolutionary analogs with a solved OspC template**

The 3D structures of the evolutionary analogs were predicted using the I-TASSER Protein Structure Prediction server [73] with the crystal structure of OspC (PDB ID: 1F1M) [45] as the template. The sequence alignment from structural superimposition of proteins was created using the UCSF Chimera (version 1.14) [74]. The overall root-mean-square-deviation (RMSD = 0.565Å) was less than 2Å and the mean alignment quality score (Q-score = 0.829) was close to 1, both indicating strong structural similarities of the evolutionary analogs with the template.

## Reference Cited

1. Allen JA, Clarke BC. Frequency dependent selection: homage to E. B. Poulton. Biol J Linn Soc. 1984;23: 15–18. doi:10.1111/j.1095-8312.1984.tb00802.x

2. Papkou A, Guzella T, Yang W, Koepper S, Pees B, Schalkowski R, et al. The genomic basis of Red Queen dynamics during rapid reciprocal host-pathogen coevolution. Proc Natl Acad Sci U S A. 2019;116: 923–928. doi:10.1073/pnas.1810402116

3. Gupta S, Maiden MC, Feavers IM, Nee S, May RM, Anderson RM. The maintenance of strain structure in populations of recombining infectious agents. Nat Med. 1996;2: 437–442.

4. Deitsch KW, Lukehart SA, Stringer JR. Common strategies for antigenic variation by bacterial, fungal and protozoan pathogens. Nat Rev Microbiol. 2009;7: 493–503. doi:10.1038/nrmicro2145

5. Ernst JD. Antigenic Variation and Immune Escape in the MTBC. Adv Exp Med Biol. 2017;1019: 171–190. doi:10.1007/978-3-319-64371-7_9

6. Pilosof S, He Q, Tiedje KE, Ruybal-Pesántez S, Day KP, Pascual M. Competition for hosts modulates vast antigenic diversity to generate persistent strain structure in Plasmodium falciparum. PLOS Biol. 2019;17: e3000336. doi:10.1371/journal.pbio.3000336

7. Ahmed Y, Tian M, Gao Y. Development of an anti-HIV vaccine eliciting broadly neutralizing antibodies. AIDS Res Ther. 2017;14: 50. doi:10.1186/s12981-017-0178-3

8. Crevar CJ, Carter DM, Lee KYJ, Ross TM. Cocktail of H5N1 COBRA HA vaccines elicit protective antibodies against H5N1 viruses from multiple clades. Hum Vaccines Immunother. 2015;11: 572–583. doi:10.1080/21645515.2015.1012013

9. Houser K, Subbarao K. Influenza vaccines: challenges and solutions. Cell Host Microbe. 2015;17: 295–300. doi:10.1016/j.chom.2015.02.012

10. Schwartz AM, Hinckley AF, Mead PS, Hook SA, Kugeler KJ. Surveillance for Lyme Disease – United States, 2008-2015. Morb Mortal Wkly Rep Surveill Summ Wash DC 2002. 2017;66: 1–12. doi:10.15585/mmwr.ss6622a1

11. Adeolu M, Gupta RS. A phylogenomic and molecular marker based proposal for the division of the genus Borrelia into two genera: the emended genus Borrelia containing only the members of the relapsing fever Borrelia, and the genus Borreliella gen. nov. containing the members of the Lyme disease Borrelia (Borrelia burgdorferi sensu lato complex). Antonie Van Leeuwenhoek. 2014;105: 1049–1072. doi:10.1007/s10482-014-0164-x

12. Margos G, Marosevic D, Cutler S, Derdakova M, Diuk-Wasser M, Emler S, et al. There is inadequate evidence to support the division of the genus Borrelia. Int J Syst Evol Microbiol. 2017;67: 1081–1084. doi:10.1099/ijsem.0.001717

13. Casjens S, Palmer N, Van Vugt R, Mun Huang W, Stevenson B, Rosa P, et al. A bacterial genome in flux: the twelve linear and nine circular extrachromosomal DNAs in an infectious isolate of the Lyme disease spirochete Borrelia burgdorferi. Mol Microbiol. 2000;35: 490–516. doi:10.1046/j.1365-2958.2000.01698.x

14. Mongodin EF, Casjens SR, Bruno JF, Xu Y, Drabek EF, Riley DR, et al. Inter- and intra-specific pan-genomes of Borrelia burgdorferi sensu lato: genome stability and adaptive radiation. BMC Genomics. 2013;14: 693. doi:10.1186/1471-2164-14-693

15. Tilly K, Bestor A, Rosa PA. Lipoprotein succession in Borrelia burgdorferi: similar but distinct roles for OspC and VlsE at different stages of mammalian infection. Mol Microbiol. 2013;89: 216–227. doi:10.1111/mmi.12271

16. Aslam B, Nisar MA, Khurshid M, Farooq Salamat MK. Immune escape strategies of Borrelia burgdorferi. Future Microbiol. 2017;12: 1219–1237. doi:10.2217/fmb-2017-0013

17. Xu Q, McShan K, Liang FT. Essential protective role attributed to the surface lipoproteins of Borrelia burgdorferi against innate defences. Mol Microbiol. 2008;69: 15–29. doi:10.1111/j.1365-2958.2008.06264.x

18. Tilly K, Krum JG, Bestor A, Jewett MW, Grimm D, Bueschel D, et al. Borrelia burgdorferi OspC protein required exclusively in a crucial early stage of mammalian infection. Infect Immun. 2006;74: 3554–3564. doi:10.1128/IAI.01950-05

19. Carrasco SE, Troxell B, Yang Y, Brandt SL, Li H, Sandusky GE, et al. Outer surface protein OspC is an antiphagocytic factor that protects Borrelia burgdorferi from phagocytosis by macrophages. Infect Immun. 2015;83: 4848–4860. doi:10.1128/IAI.01215-15

20. Önder Ö, Humphrey PT, McOmber B, Korobova F, Francella N, Greenbaum DC, et al. OspC is potent plasminogen receptor on surface of Borrelia burgdorferi. J Biol Chem. 2012;287: 16860–16868. doi:10.1074/jbc.M111.290775

21. Barbour AG, Travinsky B. Evolution and distribution of the ospC Gene, a transferable sero-type determinant of Borrelia burgdorferi. mBio. 2010;1. doi:10.1128/mBio.00153-10

22. Wilske B, Preac-Mursic V, Jauris S, Hofmann A, Pradel I, Soutschek E, et al. Immunological and molecular polymorphisms of OspC, an immunodominant major outer surface protein of Borrelia burgdorferi. Infect Immun. 1993;61: 2182–2191.

23. Bockenstedt LK, Hodzic E, Feng S, Bourrel KW, de Silva A, Montgomery RR, et al. Borrelia burgdorferi strain-specific Osp C-mediated immunity in mice. Infect Immun. 1997;65: 4661–4667.

24. Jacquet M, Durand J, Rais O, Voordouw MJ. Cross-reactive acquired immunity influences transmission success of the Lyme disease pathogen, Borrelia afzelii. Infect Genet Evol J Mol Epidemiol Evol Genet Infect Dis. 2015;36: 131–140. doi:10.1016/j.meegid.2015.09.012

25. Bhatia B, Hillman C, Carracoi V, Cheff BN, Tilly K, Rosa PA. Infection history of the blood-meal host dictates pathogenic potential of the Lyme disease spirochete within the feeding tick vector. PLOS Pathog. 2018;14: e1006959. doi:10.1371/journal.ppat.1006959

26. Wang I-N, Dykhuizen DE, Qiu W, Dunn JJ, Bosler EM, Luft BJ. Genetic Diversity of ospC in a Local Population of Borrelia burgdorferi sensu stricto. Genetics. 1999;151: 15–30.

27. Haven J, Vargas LC, Mongodin EF, Xue V, Hernandez Y, Pagan P, et al. Pervasive Recombination and Sympatric Genome Diversification Driven by Frequency-Dependent Selection in Borrelia burgdorferi, the Lyme Disease Bacterium. Genetics. 2011;189: 951–966. doi:10.1534/genetics.111.130773

28. Rannala B, Qiu WG, Dykhuizen DE. Methods for estimating gene frequencies and detecting selection in bacterial populations. Genetics. 2000;155: 499–508.

29. Qiu W-G, Dykhuizen DE, Acosta MS, Luft BJ. Geographic Uniformity of the Lyme Disease Spirochete (Borrelia burgdorferi) and Its Shared History With Tick Vector (Ixodes scapularis) in the Northeastern United States. Genetics. 2002;160: 833–849.

30. Hoen AG, Margos G, Bent SJ, Diuk-Wasser MA, Barbour A, Kurtenbach K, et al. Phylogeography of Borrelia burgdorferi in the eastern United States reflects multiple independent Lyme disease emergence events. Proc Natl Acad Sci. 2009;106: 15013–15018. doi:10.1073/pnas.0903810106

31. Di L, Wan Z, Akther S, Ying C, Larracuente A, Li L, et al. Genotyping and Quantifying Lyme Pathogen Strains by Deep Sequencing of the Outer Surface Protein C (ospC) Locus. J Clin Microbiol. 2018;56. doi:10.1128/JCM.00940-18

32. Brisson D, Dykhuizen DE. ospC Diversity in Borrelia burgdorferi Different Hosts Are Different Niches. Genetics. 2004;168: 713–722. doi:10.1534/genetics.104.028738

33. States SL, Brinkerhoff RJ, Carpi G, Steeves TK, Folsom-O’Keefe C, DeVeaux M, et al. Lyme disease risk not amplified in a species-poor vertebrate community: similar Borrelia burgdorferi tick infection prevalence and OspC genotype frequencies. Infect Genet Evol J Mol Epidemiol Evol Genet Infect Dis. 2014;27: 566–575. doi:10.1016/j.meegid.2014.04.014

34. Durand J, Herrmann C, Genné D, Sarr A, Gern L, Voordouw MJ. Multistrain Infections with Lyme Borreliosis Pathogens in the Tick Vector. Appl Environ Microbiol. 2017;83: e02552–16. doi:10.1128/AEM.02552-16

35. Izac JR, Camire AC, Earnhart CG, Embers ME, Funk RA, Breitschwerdt EB, et al. Analysis of the antigenic determinants of the OspC protein of the Lyme disease spirochetes: Evidence that the C10 motif is not immunodominant or required to elicit bactericidal antibody responses. Vaccine. 2019;37: 2401–2407. doi:10.1016/j.vaccine.2019.02.007

36. Baum E, Randall AZ, Zeller M, Barbour AG. Inferring Epitopes of a Polymorphic Antigen Amidst Broadly Cross-Reactive Antibodies Using Protein Microarrays: A Study of OspC Proteins of Borrelia burgdorferi. PLoS ONE. 2013;8: e67445. doi:10.1371/journal.pone.0067445

37. Ivanova L, Christova I, Neves V, Aroso M, Meirelles L, Brisson D, et al. Comprehensive seroprofiling of sixteen B. burgdorferi OspC: implications for Lyme disease diagnostics de-sign. Clin Immunol Orlando Fla. 2009;132: 393–400. doi:10.1016/j.clim.2009.05.017

38. Oliver LD, Earnhart CG, Virginia-Rhodes D, Theisen M, Marconi RT. Antibody profiling of canine IgG responses to the OspC protein of the Lyme disease spirochetes supports a multivalent approach in vaccine and diagnostic assay development. Vet J. 2016;218: 27–33. doi:10.1016/j.tvjl.2016.11.001

39. Earnhart CG, Marconi RT. Construction and analysis of variants of a polyvalent Lyme disease vaccine: approaches for improving the immune response to chimeric vaccinogens. Vaccine. 2007;25: 3419–3427. doi:10.1016/j.vaccine.2006.12.051

40. Durand J, Jacquet M, Paillard L, Rais O, Gern L, Voordouw MJ. Cross-Immunity and Community Structure of a Multiple-Strain Pathogen in the Tick Vector. Appl Environ Microbiol. 2015;81: 7740–7752. doi:10.1128/AEM.02296-15

41. Barbour AG, Cook VJ. Genotyping Strains of Lyme Disease Agents Directly From Ticks, Blood, or Tissue. Borrelia burgdorferi. Humana Press, New York, NY; 2018. pp. 1–11. doi:10.1007/978-1-4939-7383-5_1

42. Walter KS, Carpi G, Evans BR, Caccone A, Diuk-Wasser MA. Vectors as Epidemiological Sentinels: Patterns of Within-Tick Borrelia burgdorferi Diversity. PLOS Pathog. 2016;12: e1005759. doi:10.1371/journal.ppat.1005759

43. Qiu W-G, Schutzer SE, Bruno JF, Attie O, Xu Y, Dunn JJ, et al. Genetic exchange and plasmid transfers in Borrelia burgdorferi sensu stricto revealed by three-way genome comparisons and multilocus sequence typing. Proc Natl Acad Sci U S A. 2004;101: 14150–14155. doi:10.1073/pnas.0402745101

44. Zinder D, Bedford T, Gupta S, Pascual M. The roles of competition and mutation in shaping antigenic and genetic diversity in influenza. PLoS Pathog. 2013;9: e1003104. doi:10.1371/journal.ppat.1003104

45. Kumaran D, Eswaramoorthy S, Luft BJ, Koide S, Dunn JJ, Lawson CL, et al. Crystal structure of outer surface protein C (OspC) from the Lyme disease spirochete, Borrelia burgdorferi. EMBO J. 2001;20: 971–978. doi:10.1093/emboj/20.5.971

46. Krupka M, Masek J, Barkocziova L, Knotigova PT, Kulich P, Plockova J, et al. The Position of His-Tag in Recombinant OspC and Application of Various Adjuvants Affects the Intensity and Quality of Specific Antibody Response after Immunization of Experimental Mice. PLOS ONE. 2016;11: e0148497. doi:10.1371/journal.pone.0148497

47. Liang FT, Alvarez AL, Gu Y, Nowling JM, Ramamoorthy R, Philipp MT. An Immunodominant Conserved Region Within the Variable Domain of VlsE, the Variable Surface Antigen of Borrelia burgdorferi. J Immunol. 1999;163: 5566–5573.

48. Waddell LA, Greig J, Mascarenhas M, Harding S, Lindsay R, Ogden N. The Accuracy of Diagnostic Tests for Lyme Disease in Humans, A Systematic Review and Meta-Analysis of North American Research. PloS One. 2016;11: e0168613. doi:10.1371/journal.pone.0168613

49. CDC. Recommendations for test performance and interpretation from the Second National Conference on Serologic Diagnosis of Lyme Disease. MMWR Morb Mortal Wkly Rep. 1995;44: 590–591.

50. Branda JA, Strle K, Nigrovic LE, Lantos PM, Lepore TJ, Damle NS, et al. Evaluation of Modified 2-Tiered Serodiagnostic Testing Algorithms for Early Lyme Disease. Clin Infect Dis. 2017;64: 1074–1080. doi:10.1093/cid/cix043

51. Pegalajar-Jurado A, Schriefer ME, Welch RJ, Couturier MR, MacKenzie T, Clark RJ, et al. Evaluation of Modified Two-Tiered Testing Algorithms for Lyme Disease Laboratory Diagnosis Using Well-Characterized Serum Samples. J Clin Microbiol. 2018;56. doi:10.1128/JCM.01943-17

52. Molins CR, Sexton C, Young JW, Ashton LV, Pappert R, Beard CB, et al. Collection and characterization of samples for establishment of a serum repository for lyme disease diagnostic test development and evaluation. J Clin Microbiol. 2014;52: 3755–3762. doi:10.1128/JCM.01409-14

53. Lathrop SL, Ball R, Haber P, Mootrey GT, Braun MM, Shadomy SV, et al. Adverse event reports following vaccination for Lyme disease: December 1998-July 2000. Vaccine. 2002;20: 1603–1608.

54. Richer LM, Brisson D, Melo R, Ostfeld RS, Zeidner N, Gomes-Solecki M. Reservoir targeted vaccine against Borrelia burgdorferi: a new strategy to prevent Lyme disease transmission. J Infect Dis. 2014;209: 1972–1980. doi:10.1093/infdis/jiu005

55. Embers ME, Narasimhan S. Vaccination against Lyme disease: past, present, and future. Front Cell Infect Microbiol. 2013;3. doi:10.3389/fcimb.2013.00006

56. Earnhart CG, Buckles EL, Marconi RT. Development of an OspC-based tetravalent, recombinant, chimeric vaccinogen that elicits bactericidal antibody against diverse Lyme disease spirochete strains. Vaccine. 2007;25: 466–480. doi:10.1016/j.vaccine.2006.07.052

57. Izac JR, Marconi RT. Diversity of the Lyme Disease Spirochetes and its Influence on Immune Responses to Infection and Vaccination. Vet Clin North Am Small Anim Pract. 2019 [cited 25 Apr 2019]. doi:10.1016/j.cvsm.2019.02.007

58. Earnhart CG, Marconi RT. An octavalent lyme disease vaccine induces antibodies that recognize all incorporated OspC type-specific sequences. Hum Vaccin. 2007;3: 281–289.

59. Schuijt TJ, Hovius JW, van der Poll T, van Dam AP, Fikrig E. Lyme borreliosis vaccination: the facts, the challenge, the future. Trends Parasitol. 2011;27: 40–47. doi:10.1016/j.pt.2010.06.006

60. Chen R, Vasilakis N. Dengue — Quo tu et quo vadis? Viruses. 2011;3: 1562–1608. doi:10.3390/v3091562

61. van de Sandt CE, Bodewes R, Rimmelzwaan GF, de Vries RD. Influenza B viruses: not to be discounted. Future Microbiol. 2015;10: 1447–1465. doi:10.2217/fmb.15.65

62. Buckee CO, Recker M, Watkins ER, Gupta S. Role of stochastic processes in maintaining discrete strain structure in antigenically diverse pathogen populations. Proc Natl Acad Sci. 2011;108: 15504–15509. doi:10.1073/pnas.1102445108

63. Stamatakis A. RAxML version 8: a tool for phylogenetic analysis and post-analysis of large phylogenies. Bioinforma Oxf Engl. 2014;30: 1312–1313. doi:10.1093/bioinformatics/btu033

64. Stajich JE. An Introduction to BioPerl. Methods Mol Biol Clifton NJ. 2007;406: 535–548.

65. Fortin F-A. DEAP: Evolutionary Algorithms Made Easy. : 5.

66. Ivanova LB, Tomova A, González-Acuña D, Murúa R, Moreno CX, Hernández C, et al. Borrelia chilensis, a new member of the Borrelia burgdorferi sensu lato complex that extends the range of this genospecies in the Southern Hemisphere. Environ Microbiol. 2013;16: 1069–80. doi:10.1111/1462-2920.12310

67. Zweig MH, Campbell G. Receiver-operating characteristic (ROC) plots: a fundamental evaluation tool in clinical medicine. Clin Chem. 1993;39: 561–577.

68. Scrucca L. GA: A Package for Genetic Algorithms in R. J Stat Softw. 2013;53: 1–37. doi:10.18637/jss.v053.i04

69. MacKay DJC. Information Theory, Inference and Learning Algorithms. Cambridge University Press; 2003.

70. Hill R. A First Course in Coding Theory. Clarendon Press; 1986.

71. Chen J, Hermelin D, Sorge M. On Computing Centroids According to the $p$-Norms of Hamming Distance Vectors. ArXiv180706469 Cs. 2019 [cited 18 Nov 2020]. Available: http://arxiv.org/abs/1807.06469

72. Schindelin J, Rueden CT, Hiner MC, Eliceiri KW. The ImageJ ecosystem: An open platform for biomedical image analysis. Mol Reprod Dev. 2015;82: 518–529. doi:10.1002/mrd.22489

73. Yang J, Yan R, Roy A, Xu D, Poisson J, Zhang Y. The I-TASSER Suite: protein structure and function prediction. Nat Methods. 2015;12: 7–8. doi:10.1038/nmeth.3213

74. Goddard TD, Huang CC, Meng EC, Pettersen EF, Couch GS, Morris JH, et al. UCSF ChimeraX: Meeting modern challenges in visualization and analysis. Protein Sci Publ Protein Soc. 2018;27: 14–25. doi:10.1002/pro.3235

